# Intermittent attachments form three-dimensional cell aggregates with emergent fluid properties

**DOI:** 10.1101/2025.09.24.678186

**Authors:** Devi Prasad Panigrahi, Giulia L. Celora, Hugh Z. Ford, Robert H. Insall, Ramray Bhat, Angelika Manhart, Philip Pearce

**Affiliations:** Department of Mathematics, University College London; Mathematical Institute, University of Oxford; Institute for the Physics of Living Systems, University College London; Laboratory for Molecular Cell Biology, University College London; Center of Cell and Developmental Biology, University College London; Department of Developmental Biology and Genetics, Indian Institute of Science Bangalore; Faculty of Mathematics, University of Vienna

## Abstract

In living systems across developmental and cancer biology, populations of cells on surfaces organize themselves into aggregates that mediate function and disease. Recent experimental studies have identified that such aggregates can have emergent fluid-like properties such as surface tension, yet the physical origin of these properties is not clear. Here, we develop a minimal cell-based model in which cell-cell and cell-substrate interactions are governed by active intermittent attachments. We explain the transition of cells from a dilute population to a dense aggregate, and quantify the emergent material properties underpinning this transition. We use our model to interpret experiments on dewetting of aggregates of MDA-MDB-231 cancer cells and shape fluctuations of surface-associated OVCAR3 cell aggregates. Finally, we show how spatial heterogeneity in attachments governs collective chemotaxis of cell aggregates. Together, these results reveal how active intermittent attachments generate cell aggregates with emergent material properties, with broad implications for development and cancer.

## I. INTRODUCTION

Aggregates of motile cells perform a diverse array of physiological and pathological functions, from embryogenesis and wound healing to cancer metastasis [1–3]. Eukaryotic cell collectives can exhibit a range of complex multicellular behaviors and structures, depending on their physical interactions with the surrounding environment, from confluent tissues with long-term cell-cell adhesion [4] to motile clusters with intermittent or transient adhesion [5], which can transition from dilute populations to dense aggregates [6]. These aggregates of motile cells have been found to possess fluid-like properties that are important for numerous biological functions. For example, in intestinal epithelial development, aggregates of mesenchymal cells drive the folding of villi by forming liquid droplet-like patterns [7] (Fig. 1A(i)). Similar fluid-like properties have been observed in feeding fronts of the social amoeba *Dictyostelium discoideum*, where droplet-like patterns formed by migrating cells arise as a consequence of the emergent surface tension of the aggregate [8] (Fig. 1A(ii)). Fluid-like properties also emerge in aggregates of malignant immune cells, which exhibit an enhanced chemotactic response when migrating as a group owing to aggregate fluidity, which allows cells to exchange positions and overcome the inhibition of chemotaxis experienced by individual cells at the front of the aggregate [9]. Despite the biological relevance of the fluid-like properties of cell aggregates, the biophysical principles linking emergent aggregate properties and biological functions are not known.

**FIG. 1.**
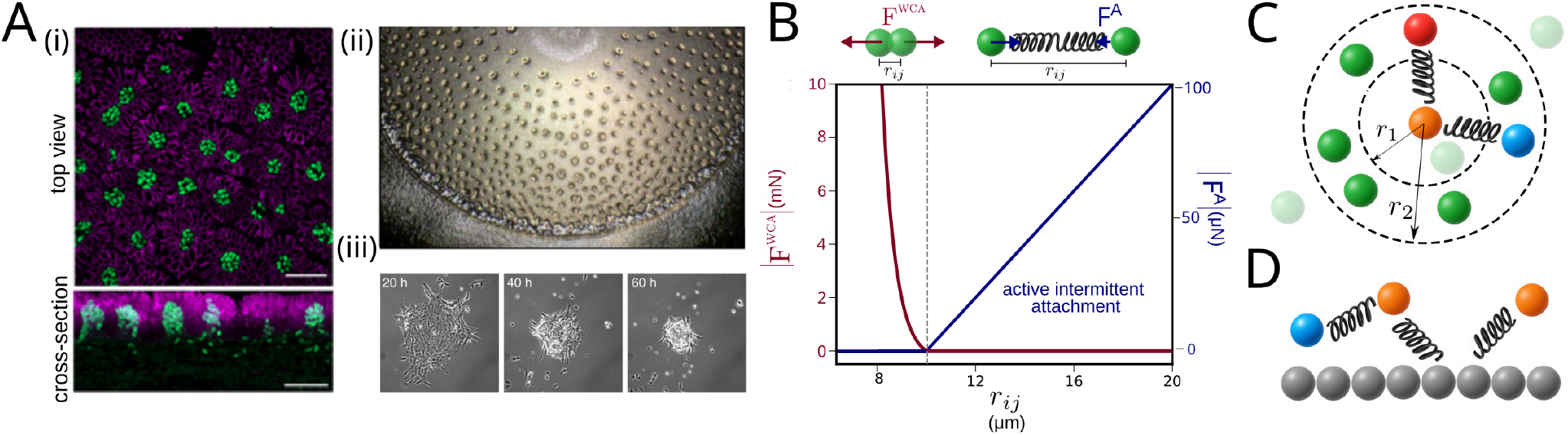
3D cell-based model with intermittent attachments. (A) Experiments showing fluid-like cell aggregates. (i) droplet-like patterns formed in the intestine of a developing embryo, where the patterns govern the folding of villi, obtained from [7] (licensed under CC BY 4.0), (ii) the pattern formed by swarms of *Dictyostelium discoideum* migrating over a bacterial lawn, obtained from [8] (licensed under CC BY 4.0), (iii) droplet-like aggregate formed by breast cancer cells (from [10], reproduced with permission from SNCSC). (B) Magnitude of the force due to WCA repulsion and intermittent active attachment between two attached cells. (C) Schematic of the cell-cell attachments showing the formation of active attachments. Colors are for labeling only – all cells behave the same. The orange cell can form an attachment to any other cell in the region between the two dashed circles with radii *r*_1_ and *r*_2_, e.g. the blue cell. It cannot attach to the faded green cells, and all other cells in the same region can also attach to the orange cell, e.g. the red colored cell forms such an attachment. See Methods for a detailed explanation, and Movie S1 for a visualization of the intermittent attachments. (D) Schematic depicting simultaneous cell-cell and cell-surface attachments. Grey spheres represent the surface and are fixed. The orange cells form attachments to the surface. The blue cell forms an attachment to the left orange cell.

In general, the formation and properties of cell aggregates depend on cell-cell and cell-substrate interactions; for example, an up-regulation of cell-cell adhesion leads to the three-dimensional (3D) aggregation of mammalian cells both *in vitro* [10] (Fig. 1A(iii)) and *in vivo* [7] (Fig. 1A(i)). Similarly, the relative strength of cell-cell and cell-substrate adhesion has been shown to regulate the spreading dynamics of cell aggregates [11, 12]. Furthermore, the classification o f different cell types within a developing embryo can be explained by the effective surface tension of aggregates [13]. Intermittent cell-cell adhesions have been hypothesized to result in emergent fluid-like properties such as viscosity and surface tension, which play a key role in the self-organization of cell aggregates [8, 14–16]. However, it is not known which physical aspects of cell-cell and cell-surface interactions control aggregate formation, three-dimensional aggregate structure, and aggregate properties.

In terms of modeling, active hydrodynamic theories have been useful in providing a macroscopic description of tissue dynamics [17], but do not typically connect emergent phenomena directly to cell behaviors. By contrast, vertex models can explain the emergence of fluid-like properties of epithelial tissues owing to mechanical interactions between confluent cells [18–20]. For example, such models have been applied to understand the role of sub-cellular structures on the emergent material properties of embryonic tissues [21]. However, since vertex models are generally applicable to confluent monolayers, they have not yet been applied to capture the emergence of three-dimensional aggregates of motile cells from di-lute populations. An alternative modeling approach is to represent cells as discrete particles with interaction forces between them. This approach has been applied to capture the emergence and dynamics of three-dimensional surface-associated bacterial aggregates through growth and physical cell-cell interactions such as steric repulsion, and attraction mediated by extracellular matrix proteins [22, 23]. More widely, particle-based models have been used to predict specific physical properties of dense aggregates, including the size and coalescence of bacterial aggregates, caused by retractable appendages such as type IV pili [15, 16]; coalescence and solid-to-fluid transitions in stem cell aggregates with intermittent attractions [24]; surface tension of surface-associated amoeboid cell aggregates with permanent cell-cell adhesions [25]; and the organization of two-dimensional cell aggregates as a function of contact inhibition of locomotion [26]. However, it is not well understood how cell interactions mediate the transition from dilute surface-associated populations to dense aggregates with emergent properties. We therefore lack a multiscale physical framework for the self-organization of eukaryotic cells into three-dimensional structures with fluid-like material properties.

Here, to bridge these gaps in biological and physical understanding, we develop a cell-based model with intermittent or transient cell-cell interactions as a coarsegrained representation of the dynamics of the actin cytoskeleton. We hypothesize that the cytoskeletal activity of the cell, along with cell-cell adhesion, can give rise to intermittent attractive forces between cells, which we model as pairwise active reciprocal interactions. We show that differential adhesion to the surface or substrate, along with intermittency in attractive forces, can explain the dewetting of cell layers to form three-dimensional aggregates. Despite the simplicity of our model, we show that it can explain several key features observed in experiments; our hypotheses can be verified in future experiments that measure how different dynamics of individual cells (e.g. between different cell lines or under different conditions) affect emergent properties of aggregates.

## II. CELL-BASED MODEL WITH INTERMITTENT ATTACHMENTS

Motivated by recent experiments showing fluid-like cell aggregates in various living systems (Fig. 1A), we aimed to understand how eukaryotic cells on substrates organize themselves into aggregates with macroscopic or emergent properties. A common feature of eukaryotic cell motility is the presence of physical interactions between cells and their surrounding environment through actin-rich protrusions on the cell membrane, such as lamellipodia or pseudopodia, that are formed along the direction of the cell’s polarity [27]. These have been observed in a diverse range of cells, including cells within developing embryos [28], neutrophils [29], fibroblasts [30], and metastatic cancer cells [31]. During motility, these protrusions form temporary attachments to features of the external environment, such as the membrane of a nearby cell or the solid substrate over which a cell is migrating [28, 32]. Cells can generate local and transient traction on the surrounding environment through attached protrusions [33] allowing them to move forward in the direction of polarization. The polarity of motile cells is dynamic, so that a protrusion can detach and retract, and a new one may form and extend in a different direction [34]. Physically, these features of eukaryotic cell motility can be considered as effectively causing cells to intermittently attach and pull themselves towards external surfaces or other cells, which feel an equal and opposite force.

To capture these features of cell motility and physical cell interactions in both dilute and dense populations of cells, we built a minimal 3D cell-based model with active intermittent or transient cell-cell and cell-surface attachments. In our model, cells are represented as rigid spheres which cannot overlap. We model this using a soft repulsive Weeks-Chandler-Andersen (WCA) potential between any two cells, so that the *i*^th^ cell experiences a force

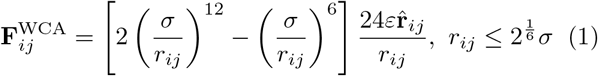

because of the presence of the *j*^th^ cell. In Eq. (1), *ε* denotes the strength of the repulsive force, **r**_*ij*_ is the inter-sphere distance, *r*_*ij*_ = |**r**_*ij*_| denotes the separation distance, 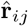 denotes the unit vector along the position vector between the two cells, and *σ* is the length scale of repulsion (corresponding to the volume exclusion experienced by the cells). When the distance between the centroids of two cells becomes less than 2^1*/*6^*σ*, they are pushed apart (Fig. 1B) to avoid overlapping. For *r*_*ij*_ *>* 2^1*/*6^*σ*, the repulsive force is zero.

In the model, we capture active intermittent attachments by introducing transient springs that form between cells and their neighbors, and pull them towards each other (Fig. 1C; see Methods for more details). Briefly, at all times, each cell is assumed to be polarized along a certain direction towards a neighboring cell within a certain distance; a temporary attachment connects the two cells. The attachment is represented as a Hookean spring with a natural length that is shorter than the current distance between the two cells. The spring remains present for a certain duration of time, sampled from a Gaussian distribution with mean *T*_*a*_ (corresponding to the timescale for persistence of cell polarity) and variance *T*_*v*_, and then disappears. The cell then instantaneously re-polarizes, and a new spring is formed with a neighbor (this may be the same neighbor as previously or a different neighbor). Therefore, at any moment in time, each cell is polarized (i.e. forms an attachment) towards exactly one neighboring cell. However, a cell might also be attached to other cells through their polarities (Fig. 1C, Movie S1). Mathematically, these interactions are captured in a transient interaction force between cells and their neighboring cells

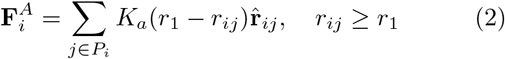

where *P*_*i*_ denotes the set of interacting cell partners of the *i*^th^ cell, *K*_*a*_ denotes the strength of cell-cell attachment, and *r*_1_ is the minimum distance between two cells at which an attachment can form (Fig. 1C). For simplicity of modeling and implementation, the surface or substrate is represented as a layer of rigid immovable spheres with the same size as the cells, to which cells can attach in the same way as other cells (Fig. 1D). To vary cell-cell and cell-surface attachments independently, the model includes a parameter *w*_*f*_ that multiplies the strength of cell-surface attachments relative to cell-cell attachments to give

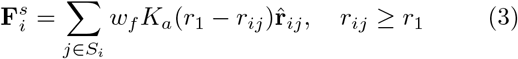

where *S*_*i*_ denotes the set of interacting surface partners of the *i*^th^ cell (the size of the set *S*_*i*_ is at most one).

Cell motility occurs at very small length scales, where the inertia of individual cells can be neglected in comparison to viscous effects. Thus, our model captures the motion of cells through the overdamped equation of motion

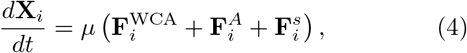

where **X**_*i*_ denotes the position of the center of the *i*^th^ cell. Cells in dense populations may form multiple simultaneous attachments to other cells [35]. However, it is known that cell motility is driven by an imbalance in forces along a particular direction arising from a dominant protrusion and associated extracellular attachments [36]; we model this temporary protrusion as a single transient spring. For simplicity, we assume that the effect of other attachments impedes the motion of the cell, and consider this effect to be implicit in the isotropic viscous response of the cell to the surrounding fluid media through the effective mobility parameter *µ*, which also captures the effect of the external fluid, and any extracellular matrix; in all simulations *µ* is assumed to be spatially uniform and constant. Eq. (4) does not contain a thermal noise term, because eukaryotic cells are too large to feel significant thermal diffusion in comparison to their active motility – stochasticity in the model arises only through the random choice of other cells when a cell forms a new attachment.

To simulate populations of cells with active intermittent attachments, we integrate Eqs. (1)-4) numerically using LAMMPS [37]. A typical simulation consists of approximately 1000 cells with periodic boundary conditions in all directions unless they are bound by a surface. Unless specified otherwise, the mean attachment time *T*_*a*_ is assumed to be equal for cell-cell and cell-surface attachments, and the default values of the various parameters in the model are chosen to be in line with experimental data (see Methods and Supplementary Materials). All quantitative data obtained from the simulations were averaged over a sufficiently large ensemble to eliminate artifacts arising from stochasticity. In general, we define a simulation as having reached a steady state when the output property of interest is roughly constant. The initial conditions were chosen depending on the system being studied (see Supplementary Materials).

## III. RESULTS AND DISCUSSION

### A. Active intermittent attachments lead to formation of 3D cell aggregates

To establish the role that active intermittent cell-cell and cell-surface attachments, which pull cells towards each other, play in self-organisation, we performed simulations of our model Eqs. (1)-(4). We found that, for certain choices of model parameters, a monolayer of surface-associated cells spontaneously self-organizes into three-dimensional (3D) aggregates that qualitatively resemble liquid droplets (Fig. 2A, Movie S2) – we refer to this as “dewetting” the surface. Cells not incorporated into aggregates remain isolated and effectively diffuse on the substrate, constituting what we term the wetting layer. We quantified this dewetting by measuring the fraction of surface area in contact with the cells (see Supplementary Materials), which decreases as the dewetted aggregates form, and plateaus at steady state (Fig. S1A). To investigate the role of intermittency in the dewetting transition, we varied the average duration *T*_*a*_ of cell attachments. When *T*_*a*_ is very small, dewetting does not occur, because the cells need to apply a force for a sufficiently long duration of time to overcome the repulsive energy barrier required for cell rearrangements and transitioning into 3D aggregates (Fig. S1B). For very large *T*_*a*_, cells are unable to rearrange because cell-cell attachments persist for too long, again precluding the formation of 3D aggregates over the timescale of our simulations, even if cell-surface attachments are intermittent (Fig. S1C,D). Single-cell tracking shows that active intermittent attachments allow cells to transition from 2D to 3D structures (Fig. 2B) and also rearrange within a 3D aggregate (Fig. S1E). These results suggest that the transient nature of active attachments allows cells to move on surfaces and form associations with other cells, causing the transition from dilute populations to dense aggregates; this mode of cell aggregation is distinct to mechanisms involving long range signaling through diffusible chemicals [38].

**FIG. 2.**
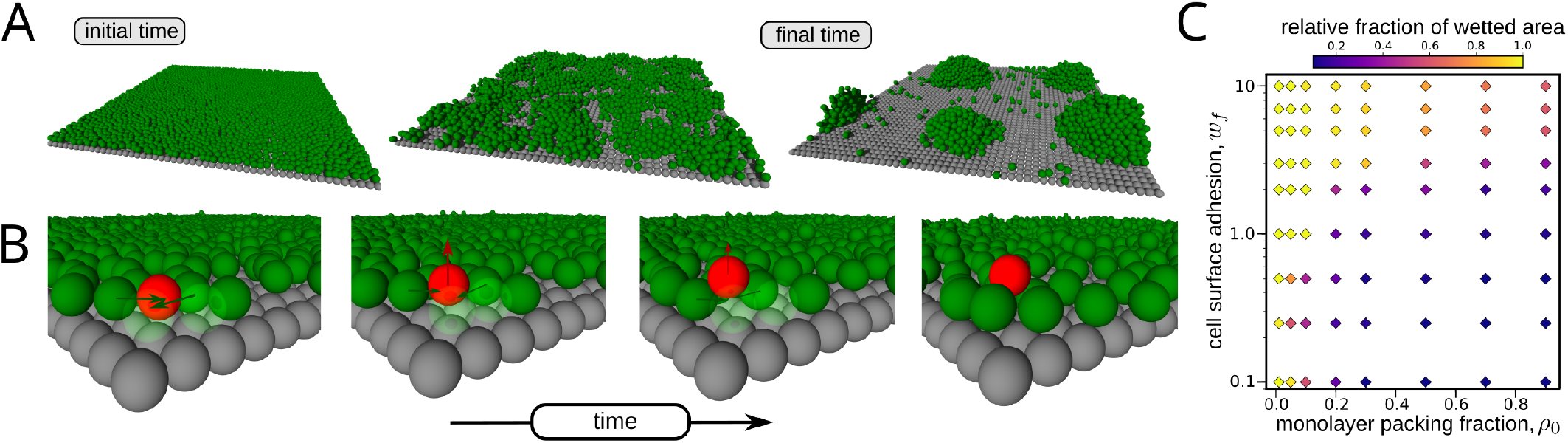
3D cell-based model with active intermittent attachments predicts formation of 3D surface associated aggregates. (A) Snapshots of a simulation showing that a monolayer of cells can spontaneously de-wet a surface, forming clusters that are nearly hemispherical in shape, closely resembling liquid droplets. The cells are represented by green spheres, and the surface by gray spheres. In this simulation *w*_*f*_ was taken to be equal to 2.0, the monolayer packing fraction in the initial condition was equal to 0.9, and the remaining parameters were taken to be equal to their default value mentioned in Table S1. (B) Single-cell tracking from a simulation showing how cell-cell interactions enable a cell to transition into the third dimension (see red cell). In the first three panels the cells in front of the red cell have been made translucent to aid visualization; arrows indicate the motion of cells. The value of *w*_*f*_ was taken to be equal to 0.25, the monolayer packing fraction in the initial condition was equal to 0.5, and the remaining parameters were same as in Table S1. (C) Variation in the fraction of the surface area wetted by the cells at steady state relative to the initial condition, as a function of differential cell-substrate adhesion, and packing fraction of the cell monolayer. In these simulations we start from a monolayer of cells and define its packing fraction *ρ*_0_ as the fraction of the total substrate area covered by cells. The colorbar represents the ratio of the surface area wetted by the cells at steady state relative to the initial condition, which is fully wetted. When the relative fraction of wetted area is close to one, the cells do not undergo any significant dewetting, and a lower value indicates greater dewetting.

To understand more broadly how cell-cell attachment strength, cell-surface attachment strength and cell density affect self-organisation in our model, we generated a regime diagram of dewetting. To do so, we performed simulations of a cell monolayer covering an initial area *ρ*_0_ with differential cell-surface vs cell-cell attachment strength *w*_*f*_ (see Eq. 3), and quantified the final area covered by aggregates, relative to the initial area *ρ*_0_ (Fig. 2C). We found that an increase in *w*_*f*_ leads to an increase in the fraction of the wetted area, irrespective of the number of cells in the monolayer (represented by *ρ*_0_), as long as there is a sufficiently large number of cells to allow enough cell-cell contacts. This shows that the transition into 3D structures and the resulting dewetting phenomenon are governed by local cell attachments. Taken together, these results show that this active system – in which cell motion is generated only by transient cell-cell and cell-surface interactions, and not by Brownian diffusion – bears qualitative similarities with passive fluids, in which wetting is governed by the relative strength of intermolecular attraction between the fluid molecules, and between the fluid and the substrate.

### B. Emergent fluid-like properties of cell aggregates

To quantify the link between active intermittent attachments and the observed fluid-like properties of surface-associated cell aggregates, we performed idealized simulations to extract the properties of cell aggregates. First, we performed simulations of isolated aggregates not associated to a surface, to show that active cell-cell attachments minimize aggregate surface area (Fig. S2A, Movie S2); this indicates the presence of an emergent surface tension. Then we measured the aggregate surface tension by simulating aggregates squeezed between two fixed plates that are non-adherent to the cells. Owing to emergent surface tension, the aggregate’s preferred shape is spherical; the confinement therefore induces a net outward force *F* on the plates. This force is related to the emergent surface tension of the aggregate Σ [25, 39]; specifically:

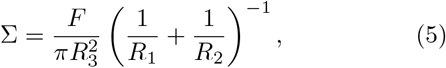

where *R*_1_, *R*_2_ and *R*_3_ are respectively the two principal radii of curvature of the aggregate and the radius formed by the contact area of the aggregate with the plates (Fig. 3A). We estimated *F, R*_1_, *R*_2_, and *R*_3_ computationally, and our simulations predict that a higher attachment strength increases emergent surface tension (Fig. 3B), in line with results from passive systems [40]. We found that surface tension varies non-monotonically with attachment time (Fig. 3C). In general, these simulations predict that active intermittent attachments generate a quantifiable surface tension in cell aggregates that depends on the strength and intermittency of cell-cell attachments in a non-trivial manner, a prediction that is testable in future experiments with a similar protocol.

**FIG. 3.**
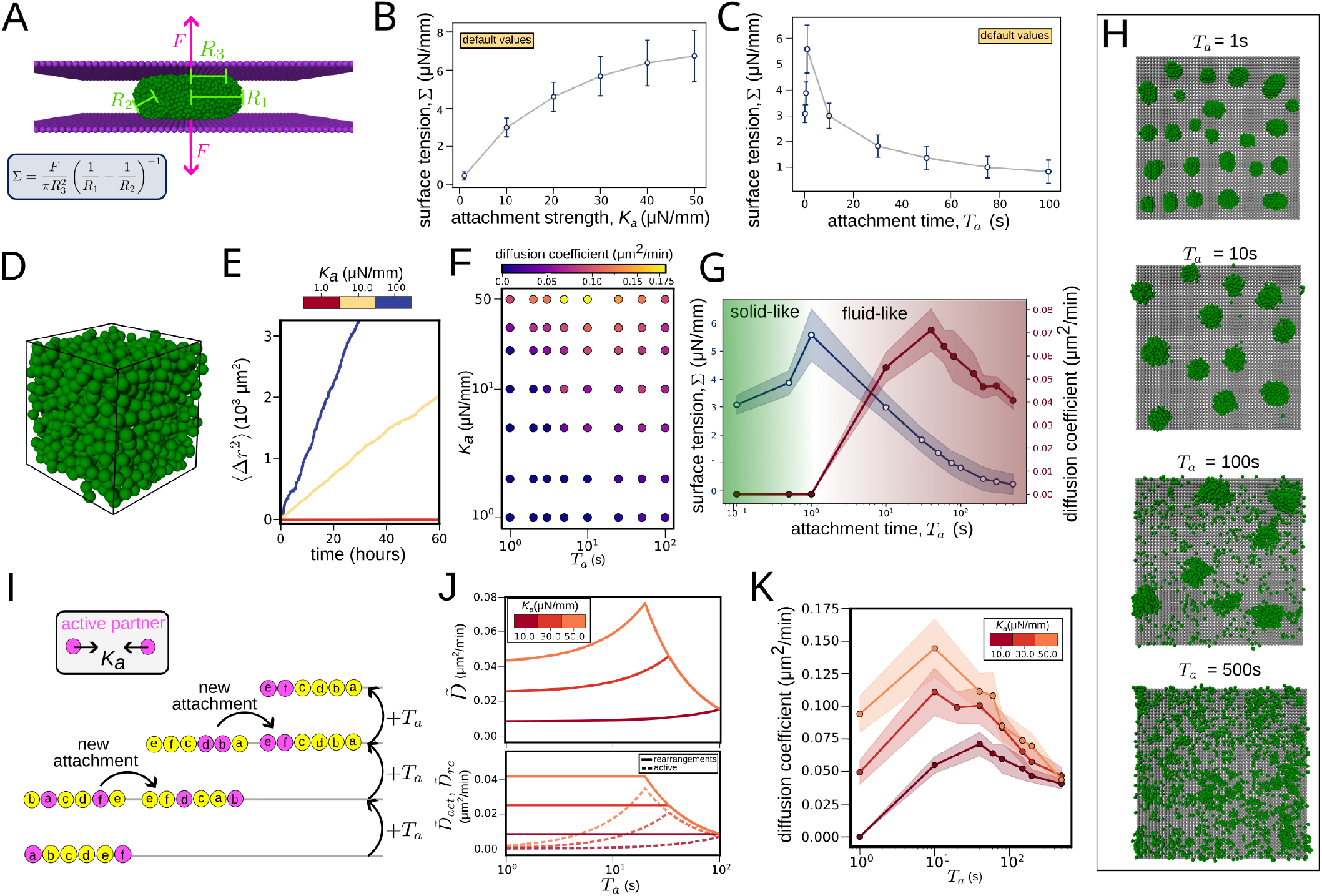
Active intermittent attachments generate quantifiable emergent surface tension and fluidity of cell aggregates. (A) Schematic of the parallel plate compression test, where the two walls are assumed to be non-adherent to the cells, unlike the simulations in previous sections. (B,C) Variation in emergent surface tension with attachment strength *K*_*a*_ (B) and attachment time *T*_*a*_ (C), for the default parameter set mentioned Table S1. (D) Schematic of the simulation used for quantifying the emergent fluidity. The black lines denote periodic boundaries. (E) Mean square displacement vs. time for different values of attachment strength when the attachment time *T*_*a*_ = 10s. (F) Variation in emergent diffusion coefficient for different values of attachment strength and attachment time. The colorbar denotes the value of the diffusion coefficient in the units of *µ*m^2^*/*min. (G) Comparison of the variation of emergent surface tension and diffusivity as a function of the attachment time when all the other parameters are same as their default values. (H) Snapshots of a cell aggregate undergoing dewetting, starting from a monolayer with a packing fraction of 0.5, for different values of *T*_*a*_. (I) Schematic of the simplified one-dimensional model used for gaining insight into the emergent fluidity. We consider a simplified 1D model where cells actively attract one another, and other cells that lie in their path undergo rearrangements. More details on the model can be found in the Supplementary Materials. (J) Emergent diffusion coefficient from the reduced 1D model. The top panel depicts the total diffusion coefficient; the bottom panel shows the contributions from active attachments and associated rearrangements. (K) Emergent diffusion coefficient in simulations vs. attachment time *T*_*a*_, for different values of attachment strength *K*_*a*_.

To quantify emergent fluidity of the aggregate arising from active intermittent cell-cell attachments, we performed simulations of unconfined cell aggregates with periodic boundary conditions (Fig. 3D), and extracted the mean-squared displacement (MSD) of cells from their initial locations. The MSD was found to increase approximately linearly with time (Fig. 3E), suggesting an emergent diffusion of cells caused by intermittent cell-cell attachments. We therefore quantified the fluidity of the cell aggregate by extracting the effective diffusion coefficient of the cells *D* [41] by fitting a straight line to the MSD vs time plot using the relationship

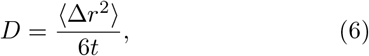

where Δ*r* denotes the displacement of a cell, *t* indicates time and ⟨·⟩denotes an average across all cells within the aggregate. Our results show that the diffusion coefficient *D* monotonically increases with the attachment strength *K*_*a*_, but has a non-monotonic dependence on the attachment time *T*_*a*_ (Fig. 3F); initially, *D* increases with increasing *T*_*a*_, but it decreases with *T*_*a*_ when *T*_*a*_ is above a critical value. Overall, our simulations predict that intermittent cell-cell attachments generate a quantifiable emergent fluidity in cell aggregates, which is controlled by the strength and intermittency of the attachments.

Physically, the aggregate-level material properties that we have extracted – surface tension and fluidity – depend on the ability of cells to rearrange their positions within an aggregate. To explore this connection, we considered how the material properties are governed by cell motility caused by active attachments. A dimensional analysis of the model reveals the key non-dimensional parameter that varies between our simulations: the persistence number *P* (see Supplementary Information), defined as

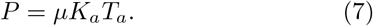

This represents the ratio between the timescale of cell–cell attachment *T*_*a*_, and the time required for a cell to move across a characteristic attachment length, 1 /*µK*_*a*_. When we plot the non-dimensional surface tension Σ*/K*_*a*_ against the persistence number, we find that our simulations collapse onto a single curve (Fig. S2B), demonstrating that the emergent surface tension is determined solely by the active nature of the intermittent attachments (when volume exclusion and attachment range are fixed). Similarly, the non-dimensional emergent diffusivity in aggregates collapses when plotted against persistence number (Supplementary Material; Fig. S2D). Persistence of cell motility explains our dimensional results as follows. In terms of fluidity, very short attachment times preclude rearrangements, because the cells do not acquire sufficient energy to push apart their neighbors and induce rearrangements (Fig. S2E). By contrast, when the attachment time *T*_*a*_ is very long, the attachments last long enough for active cell partners to come together and then remain attached while stationary, decreasing average cell speed (Fig. S2C); this explains the decrease in fluidity for very high values of *T*_*a*_. For intermediate values of *T*_*a*_, cell rearrangements occur regularly and fluidity is maximised (Fig. 3G). Our results for surface tension show a qualitatively similar dependence on attachment time, although we find that surface tension can still be present – and takes its maximal value – in solid-like aggregates with minimal cellular rearrangements (Fig. 3G). Overall, these results show that the aggregate-level properties arising from active intermittent attachments can be both solid-like and fluid-like, and the intermittency in active cell-cell attachments governs the transition between the two.

Finally, to understand the implications of these emergent material properties during the formation of cell aggregates, we performed further monolayer dewetting simulations for a range of values of *T*_*a*_ (Fig. 3H). At steady state, for a small value of *T*_*a*_, associated with a solid-like aggregate, we found that all the cells form clusters and no cells remain in the wetting layer; the clusters behave as rigid bodies (Movie S3). By contrast, for higher values of *T*_*a*_ within the fluid-like regime, we observed a finite number of cells populating the wetting layer; the number of cells in the wetting layer increases with an increase in *T*_*a*_ (Fig. 3H). For a very large value of *T*_*a*_, we observed that the aggregation of cells is arrested; this is in line with the very low values of surface tension measured for high *T*_*a*_ in our idealized simulations (Fig. 3A,G). Taken together, our results show how the active nature of inter-mittent attachments governs the material properties of cell aggregates, and how these properties are implicated in the formation of 3D surface-associated cell aggregates.

### C. Emergent fluidity arises due to a combination of active cell-cell attachments and cell rearrangements

We expect emergent diffusion to arise from a combination of “active” movement from attachments and cell rearrangements caused by the active movement of other cells (Fig. S2E). To explore in detail the importance of each type of motion, we built a simplified 1D on-lattice model in which cells occupy all lattice points and move via intermittent active attraction to other cells, while displacing the cells between them (Fig. 3I, Supplementary Materials). Specifically, in the 1D model, to form a new attachment a randomly selected cell chooses an interaction partner at random from cells within a fixed distance on either side. During attachment, the two cells move towards one another by one cell diameter each timestep; an attachment lasts for *T*_*a*_ time steps or until the cells occupy spaces next to each other, whichever is smaller. As the two attached cells move towards each other, each time one of them takes a step, which would cause them to occupy the same position as another cell, they instead switch position with that cell – this mimics displacement of cells because of volume exclusion. Only one attachment between two cells is present at any one time in the system. The simplified 1D geometry allows us to obtain an analytical expression for the emergent diffusivity 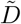 (see Supplementary Materials) which can be decomposed into an active part 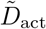, arising from the active attachment of a cell, and an accompanying rearrangement part 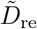 that arises from rearrangements due to the active motion of other cells,

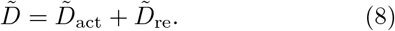

We find that

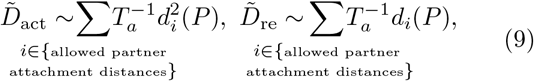

where *d*_*i*_ is the typical distance (in cell diameters) traveled during an active attachment between cells that are initially *i* cell diameters apart (see Supplementary Materials) and *P* is the persistence number (see Eq. 7). Surprisingly, this implies that, in this simplified model, for our typical parameter choices, the cell rearrangements contribute more to the emergent diffusion coefficient than active motion caused by attachments (Fig. 3J). However, as the attachment range increases, active motion can have a dominant contribution to 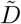 (Fig. S2F).

Our 1D model reveals in more detail the role of cell persistence on emergent aggregate fluidity. The typical distance traveled during an attachment *d*_*i*_ (for *i >* 1, i.e. non-adjacent cells) increases with the persistence number *P*, saturating at large *P* (see Supplementary Materials). Owing to the pre-factor 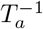 in Eq. 9, this leads to a non-monotonic variation of both 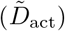, and 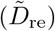 with *T*_*a*_ (Fig. 3J), thereby explaining the non-monotonic variation in emergent fluidity with attachment time, as seen in the simulations (Fig. 3K). Overall, these results demonstrate that both active attachment-induced motion and cell rearrangements are required for emergent tissue fluidity – each type of motion enables the other, and each contributes significantly to the overall emergent diffusivity of cells.

### D. Differential strength of cell-surface active intermittent attachments explains dewetting of aggregates

Once the basis for emergent fluid-like properties of cell aggregates was established, we used our model to interpret recent experimental results [10] on dewetting of aggregates of MDA-MB-231 breast cancer cells. In these experiments, it was shown that a cell monolayer can dewet a surface upon increase in cell-cell adhesion due to induction of E-Cadherin which increases the strength of cell-cell interactions while keeping the cell-substrate interactions unaltered. In Fig. 4A, we show that our model can capture the variation in wetted area with time, starting from the onset of dewetting. To understand the three-dimensional nature of this process, we consider a side view of the cell aggregate during dewetting and show that the increase in height of the aggregate with time during dewetting is also captured by our simulations (Fig. 4B).

**FIG. 4.**
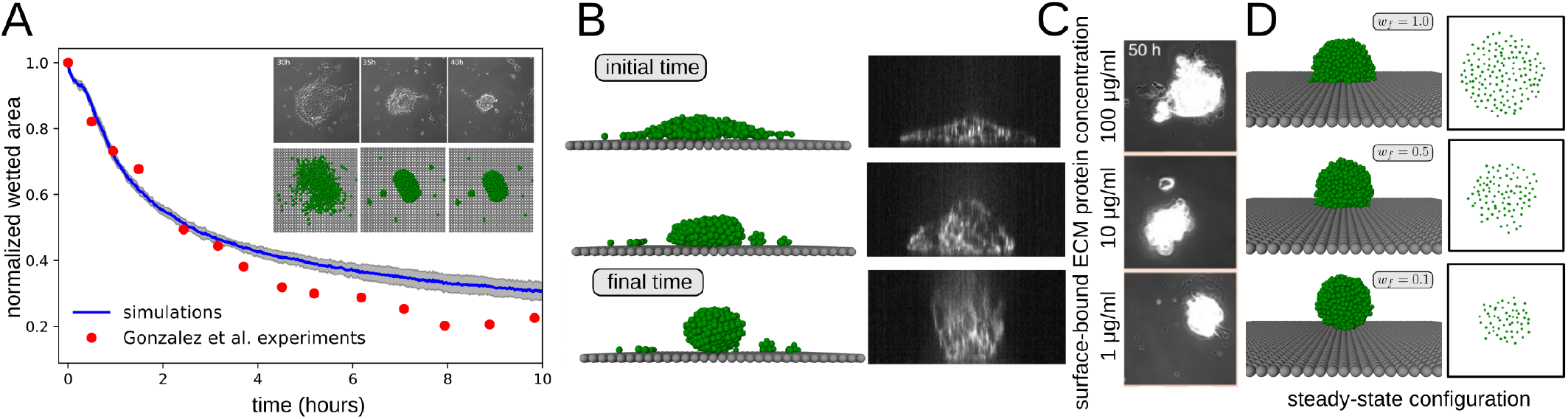
Active intermittent attachments captures the dewetting of cell aggregates. (A) Variation of the wetted area of an aggregate over time, obtained by averaging over 10 different simulations starting from the same initial condition, and experiments on aggregates of breast cancer cells (data obtained from [10]). The wetted area is normalized at all time points by its value at the onset of dewetting. Inset: top views of the aggregate from both simulations and experiments. (B) A comparison of the side view of dewetting, from both experiments and simulations. For the simulations shown in panels A and B, *w*_*f*_ = 0.01, *K*_*a*_ = 9*µN/*mm, *µ* = 5 m · *N* ^−1^ · *s*^−1^, and the values of all other parameters is same as their default values mentioned in Table S1. (C) In the experiments, a decrease in concentration of a surface-bound ECM protein (from top row to bottom row) leads to a decrease in wetted area. (D) Simulations showing the 3D cell aggregate (left), and the corresponding contact area (right) for different values of cell-surface adhesion strength *w*_*f*_ in Eq. (3). The contact areas in C and the right-most column of D, representing the cell aggregate at steady state, appear visually similar because the microscopy imaging predominantly captures the cell-substrate interface. Experimental images in this figure have been obtained from [10], and are reproduced with permission from SNCSC.

Next, we used our model to understand how the variation in relative strength of cell-cell and cell-substrate interactions can affect the steady-state wettability of cell aggregates. It was shown that a decrease in the concentration of surface-bound collagen, which is an extracellular matrix (ECM) protein, leads to a decrease in wettability (Fig. 4C) [10]. To qualitatively explain these experimental observations, we performed simulations of a single aggregate and modelled the change in collagen concentration as an effective decrease in cell-surface attachment strength by modifying the parameter *w*_*f*_ in Eq. 3. We found that a decrease in cell-surface attachment strength via *w*_*f*_ causes a reduction in wetted area of the aggregate at steady state (Fig. 4D, 2C), in line with the experimental observations (Fig. 4C) and recent results on spreading of cell aggregates [42, 43]. Taken together, our results suggest that the relative strength of active intermittent cell-cell and cell-surface attachments can explain the dewetting of cell populations and the ability of aggregates to transition into 3D structures.

### E. Strength of cell-cell and cell-surface attachments determine shape fluctuations of 3D cell aggregates

In recent experiments comparing aggregates of ovarian cancer OVCAR-3 cells to aggregates of chemotherapeutic drug-resistant OVCAR-3 cells, it has been observed that the resistant cells spread more effectively on a substrate compared to the drug-sensitive cells [44]. Drug-resistant cells were also observed to exhibit increased single-cell motility, measured via trans-well migration assay, with implications for invasiveness *in vivo*. Furthermore, aggregates of drug-resistant cells were measured to be much less circular and to exhibit more pronounced shape fluctuations than aggregates from the parental drug-sensitive cell line (Fig 5A). At the cell level, drug-resistant cells were found to exhibit increased expression of both E-Cadherin and Fibronectin (Fig. 5B), which mediate cell-cell and cell-substrate adhesion, respectively, indicating a combined increase in both cell-cell and cell-substrate adhesion. We aimed to explore a potential connection between these cell- and aggregate-level observations using our model.

**FIG. 5.**
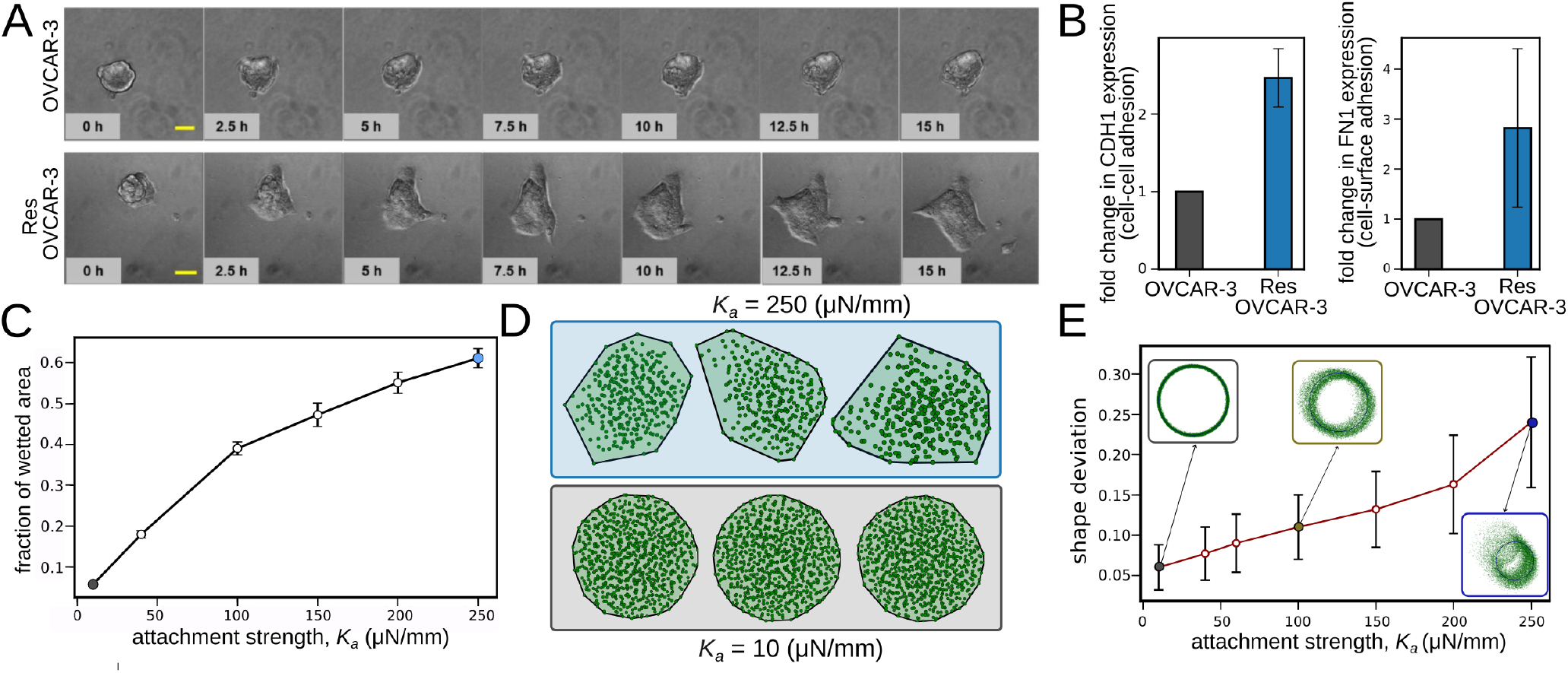
Active intermittent attachments mediate wetted area and shape fluctuations of cell aggregates on surfaces. (A) Time-lapse imaging of wetting of an aggregate of OVCAR-3 ovarian cancer cells (top), and drug-resistant OVCAR-3 ovarian cancer cells (bottom), on a substrate coated with collagen-I. Comparison of the top and bottom rows reveals that aggregates of drug-resistant OVCAR-3 cancer cells exhibit greater surface wettability and shape fluctuations (obtained from [44], licensed under CC BY-NC-ND 4.0). (B) Experimental measurements of the expression levels of E-Cadherin (left) and Fibronectin (right), which govern, respectively, cell-cell and cell-surface adhesion, reveal both proteins are upregulated in drug-resistant cancer cells. Data have been obtained from [44], licensed under CC BY-NC-ND 4.0. (C) Variation in the fraction of the surface area wetted at steady state with the attachment strength *K*_*a*_ when *w*_*f*_ = 1. An increase in *K*_*a*_ leads to a simultaneous increase in both cell-cell and cell-surface attachment strengths. (D) Snapshots of the top view of a cell aggregate at steady state (i.e. when aggregate-surface contact area remains roughly constant with time), for two different values of *K*_*a*_. The black line denotes the approximate boundary of the aggregate, included to highlight the shape fluctuations. Cells outside the aggregate are not plotted. A 3D rendering of these simulations is shown in Fig. S3C. (E) Variation in shape deviation with attachment strength *K*_*a*_for the simulations corresponding to C. The insets show the points (in green) along the boundary of the aggregate (i.e. on the convex hull; see Supplementary Materials) at different instances of time, after it has attained a steady state; we also show the time-averaged position of the boundary (blue line). A complete list of model parameters other than the ones mentioned here can be found in the Supplementary Materials (Table S1).

To investigate the effect of increasing both cell-cell and cell-surface interactions in our model, we performed simulations for different values of *K*_*a*_ in Eq. (2), which determines the strength of both types of attachment (we assumed that each attachment has the same strength, *w*_*f*_ = 1). We explored whether this was sufficient to qualitatively recapitulate experimental observations. First, to probe single-cell dynamics, we assessed the motility of isolated cells, finding that increasing *K*_*a*_ enhances motility (Fig. S3A), in line with experiments [44] (Fig. S3B). To probe the collective behavior, we then simulated 3D aggregates. As before, we assume the aggregate is at steady state when the contact area of the aggregate with the surface remains roughly constant with time. We found that an increase in *K*_*a*_ leads to a monotonic increase in the fraction of wetted area at steady state (Fig. 5C, Fig. S3C). To analyze shape fluctuations in our simulations, we extracted the convex hull of cell centroids within an aggregate, when projected onto a plane parallel to the surface, and quantified its deviation from a circle (see Supplementary Materials). We defined a cell as belonging to the aggregate if they were not in the wetting layer near the surface (see Supplementary Materials). We found that for higher values of *K*_*a*_, the aggregates contain fewer cells due to enhanced wetting, and their shape both fluctuates (Fig. 5D) and deviates (Fig. 5E) from a circle by a greater amount. These fluctuations lead to the extension of multicellular protrusions along the aggregate boundary (Movie S4), in agreement with the experiments (Fig. 5A). These shape fluctuations are driven by the increase in cell attachment strength – for example, increasing the cell-surface differential adhesion parameter *w*_*f*_ increases wetting without strongly affecting the magnitude of shape fluctuations (Fig. S3D). These results suggest that, although chemotherapeutic resistance of cancer cells might be expected to induce a wide range of changes in cell behavior, changes to cell attachments are sufficient to explain observed changes to overall aggregate morphology. In general, our simulations predict that the overall strength of intermittent cell-cell and cell-surface attachments increases shape fluctuations and wetting of surface-associated cell aggregates.

Similarly to the emergent material properties explored previously, aggregate wetting and fluctuations can be explained in terms of the persistence of cell motility caused by active attachments. Enhanced persistence of cell motion causes cells near the aggregate boundary that are polarized towards the surface in a direction external to the aggregate to travel further along the surface before either returning to the aggregate or escaping entirely. This suggests increased persistence enhances both wetting and shape fluctuations, as shown in Figs. S3E,F. Furthermore, experiments in conditions that promote monolayer formation, rather than 3D aggregation, show that monolayers of drug-resistant OVCAR-3 cells – which we expect to be more persistent based on their promoted cell-cell and cell-surface adhesion – exhibit greater fluidity than OVCAR-3 monolayers (Fig. S3G), consistent with our results on increased emergent diffusivity with increase in persistence (Fig. S2D). Taken together, these results suggest that the strength and intermittency of cell-cell and cell-surface attachments govern biologically relevant properties of cell aggregates through the active motion of individual cells.

### F. Bias in active intermittent attachments generates collective migration of 3D cell aggregates

Collective cell migration is a fundamental biological phenomenon with physiological relevance in disease [45] and development [46]. Recently, aggregates or swarms of *Dictyostelium discoideum* cells have been found to collectively migrate towards nutrients because of chemotactic bias, while displaying extensive cell rearrangements and pattern formation caused by periodic splitting of the swarm front as it migrates [8] (Figs. 1A(ii) and 6A). Similar cell rearrangements have been observed in the collective chemotaxis of malignant immune cells [9]. Motivated by these experiments, we aimed to explore whether intermittent attachments are sufficient to explain collective chemotaxis of cell aggregates, pattern formation caused by aggregate splitting, and cell rearrangements within aggregates during migration.

First, we explored whether biased active intermittent attachments are sufficient to explain collective migration of aggregates. We performed simulations of aggregates with bias in the direction in which cells’ active intermittent attachments form, determined by a self-generated chemoattractant gradient (see Supplementary Materials, Fig. S5). A well characterized form of robust and longrange chemotaxis involves the localized consumption of chemoattractant by cells, giving rise to a self-generated gradient in chemoattractant concentration which replenishes as cells move towards the chemoattractant [47]. As a simple representation of this process, we considered a one-dimensional field of diffusive chemoattractant that is consumed locally by each cell (see Fig. 6B,C, Supplementary Materials). To represent bias in the model, we allowed for preferential attachment to neighbors in the direction of increasing chemoattractant, with a probability that increases with the strength of the gradient in receptor occupancy (with a plateau at high occupancy; see Supplementary Materials). We found that this was sufficient for long-distance migration of aggregates, which follow a chemoattractant concentration gradient that travels with the aggregate (Fig. 6B). During migration, the bias in the direction of attachment is non-uniform within the aggregate – bias is higher towards the leading edge of the aggregate, where cells experience a higher gradient in chemoattractant concentration because of chemoattractant consumption by cells within the aggregate (Fig. 6B). A higher bias in the leading edge of the aggregate might be expected to cause the aggregate to spread out over time, with leading cells traveling faster. Despite this, we find that for certain values of the parameters, the aggregates maintain an approximately steady shape and speed during migration (Fig. S4A); by contrast, under conditions that favor wetting, aggregates do not remain coherent during migration (Fig. S4B). Furthermore, we find that the migration speed of aggregates is generally similar to the speed at which they would migrate if the attachment bias of all cells were imposed to be the same as the average self-generated bias (see Fig. 6D). In line with experiments and continuum modeling [8], this shows the implication of emergent surface tension on holding together a migrating aggregate which would otherwise spread out as each cell moves according to its own chemotactic bias [48].

**FIG. 6.**
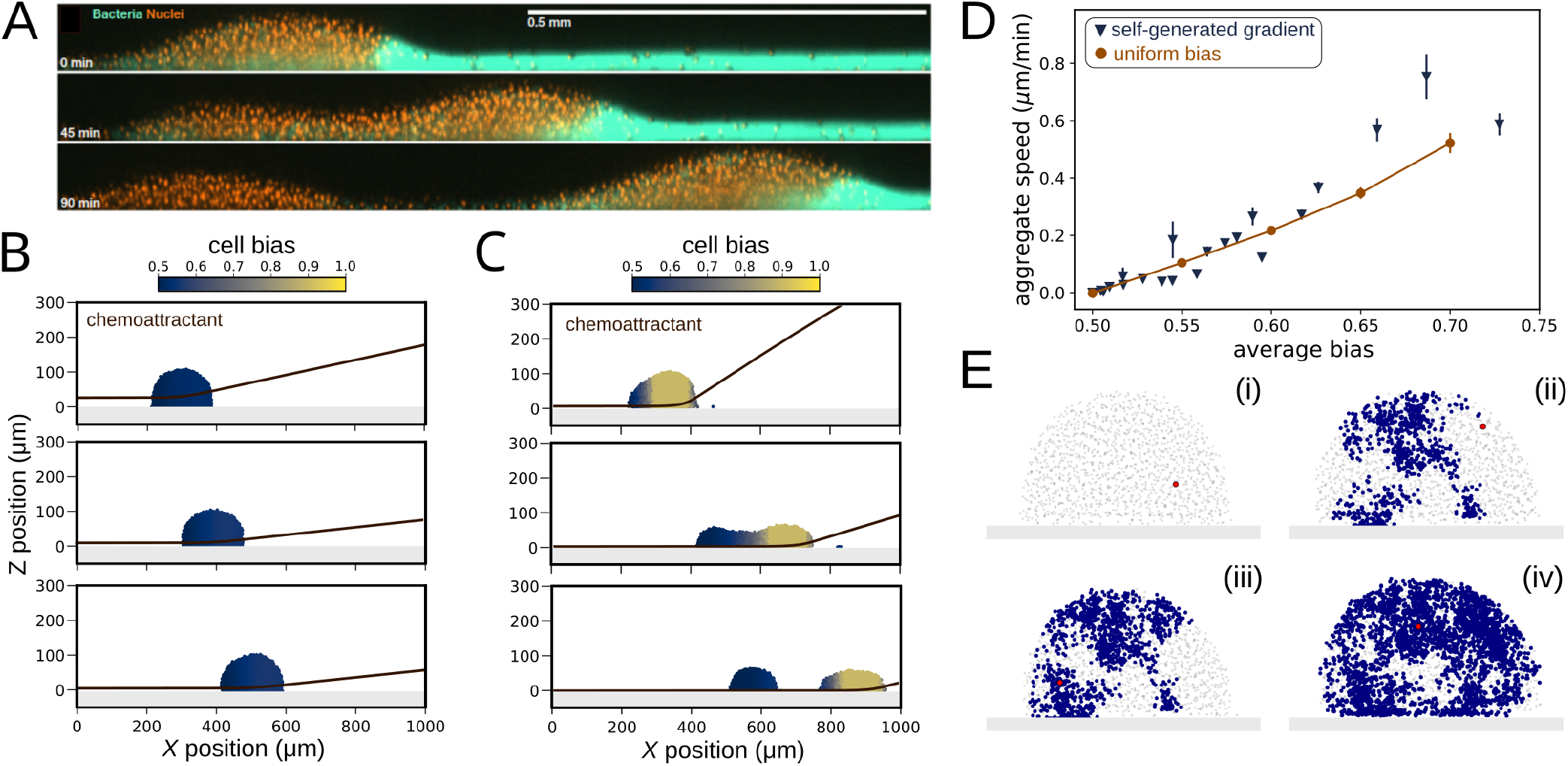
Active intermittent attachments with bias cause migration of cell aggregates. (A) Snapshots from experiments on aggregates of *Dictyostelium discoideum* undergoing chemotaxis to bacteria (obtained from [8], licensed under CC BY 4.0). The orange fluorescent colored dots denote the position of the cell nuclei, and the green fluorescent color denotes the bacteria. (B) Migration of a cell aggregate with surface tension owing to self-generated chemotaxis and biased intermittent attachments. The color denotes bias experienced by single cells, and time increases from top to bottom. Here, the position of the cells is overlaid with the chemoattractant concentration profile, which is appropriately scaled, in order to visualize the collective chemotaxis. Cell bias depends on the gradient in receptor occupancy (see Supplementary Materials); note that the slight reduction in bias at the front of the aggregate is caused by saturation of receptor occupancy. The surface is illustrated by the gray shaded area, and bias affects both cell-cell and cell-surface interactions. The aggregate stays together despite the spatial variation in bias, indicating an emergent surface tension. By contrast, under conditions that favor wetting, the aggregate spreads out (Fig. S4B). Further details on the boundary conditions for the chemoattractant concentration field, model parameters (Table S1), and mathematical implementation are in the Supplementary Materials. (C) Migration and splitting of an aggregate of cells because of emergent surface tension from intermittent attachments; splitting has qualitative similarities to experiments in panel (A). In the model, the rate of degradation of the chemoattractant is higher in C as compared to B, which causes splitting. Other factors that cause splitting include increased aggregate size and chemotactic sensitivity; see Fig. S4. (D) Comparison of the migration speed of aggregates in self-generated chemotaxis (similar to panels B and C), and simulations where all the cells in the aggregate are fixed to have the same value of bias (i.e. uniform bias model). The migration speeds of the aggregates in self-generated chemotaxis are similar to those of uniformly biased aggregates with the same average bias. (E) Intermittent attachments generate an emergent tissue fluidity in migrating aggregates. Footprint of a single particle within an aggregate (time increasing from (i) - (iv)), for the simulation corresponding to (B). The red colored dot denotes the current position of the cell, and the blue color denotes the positions occupied by the cell in all preceding instances of time.

Second, to investigate how fluid-like properties emerge and mediate pattern formation in migrating aggregates, we aimed to reproduce previously observed splitting of swarms during migration [8]. We found that an enhanced heterogeneity of bias within a migrating aggregate can lead to splitting, where a small aggregate of non-biased cells is left behind an aggregate that continues to migrate at reduced size (Fig. 6C, Movie S6). For a given parameter set, there exists a critical size of the aggregate beyond which it splits (Fig. S4C,D). This can explain the repeated splitting events seen in experiments and continuum modeling of active droplets [8], where cell proliferation gradually increases the size of the aggregate to its critical size after each splitting event. An enhanced spatial heterogeneity in bias can also be induced by increasing the rate of chemoattractant degradation or the chemotactic sensitivity of cells (Figs. S4C,E). To characterize these splitting events, we considered the total aggregate span which we define to be the difference between the maximum and minimum *x* coordinates of cells that are not in the wetting layer. After a splitting event, this parameter increases steadily with time as the gap between the two parts of the aggregate keeps increasing with time (see Fig. 6C); the aggregate span can therefore be used to classify whether an aggregate splits. By performing simulations for different values of initial aggregate size, chemotactic bias and consumption rate, we found that a sufficiently large value of any one of these parameters can cause splitting (Fig. S4F-H). Together, these results show that emergent aggregate properties arising from active intermittent attachments between cells can explain pattern formation caused by splitting of migrating aggregates during collective chemotaxis.

Finally, we investigated whether active intermittent attachments are sufficient to explain cell rearrangements during aggregate migration. In general, experiments reveal that cells move around within migrating aggregates over timescales much faster than cell thermal diffusivity [8, 9, 49]. This process is observed across many tissue types and is referred to as “tissue fluidity” [50, 51]. Specifically, in *D. discoideum*, it has been found that this fluidity prevents the occurrence of starvation-induced differentiation within the population during migration [52]. In aggregates of cancerous immune cells, fluidity contributes to the efficiency of collective chemotaxis by the regular replacement of chemotactically impaired cells that have been exposed to a high concentration of chemoattractant at the front of the aggregate [9]. Our simulations show that active intermittent attachments generate fluidity within a migrating aggregate, so that individual cells are able to move between the front of the aggregate, where their polarity is highly biased and nutrients are expected to be plentiful, and the back of the aggregate, where their bias is low and nutrients are expected to be scarce (Fig. 6E, Fig. S4I); we do not separate nutrients and chemoattractants in our modeling, but their concentrations are often correlated [8]. When this fluidity is absent – for example, for very small values of the attachment time *T*_*a*_, for which cell rearrangements are unable to occur (Fig. S1B) – we find that a solid-like aggregate migrates in the direction of bias (Movie S5). Overall, these results suggest that biased active intermittent attachments generate an emergent cell motion that causes the rearrangement of cells within migrating aggregates, with significant consequences for biological outcomes in individual cells.

## IV. CONCLUSION

Here, we have used a minimal cell-based model to demonstrate how active intermittent cell-cell and cellsurface attachments cause the self-organization of cell populations into 3D aggregates with emergent fluid properties. Our approach – an off-lattice cell-based model in which cells move via active intermittent interaction forces – allowed us to capture how dilute, surface-associated cell layers transition to dense 3D aggregates with extensive cell rearrangements. We have connected the formation and properties of aggregates directly to cell-cell interactions by explicitly modelling the transience of such interactions (as opposed to modelling cell motion via an effective Brownian diffusion). Specifically, by extracting and analyzing the motion of all cells within aggregates, we have established and quantified how intermittent attachments generate emergent fluid-like properties including surface tension, wettability and shape fluctuation in aggregates. Furthermore, we have found that within aggregates, cells undergo random motion owing to an effective diffusivity (which is much larger than thermal diffusivity) due to active motion and associated cell rearrangements enabled by intermittency in active cell-cell attachments. Broadly, we have found that emergent fluid-like properties of aggregates are governed by the persistence of cell motion, which here is governed by the transient cell-cell and cell-surface attachments. By comparing our results qualitatively with experimental data, we have shown that our model is able to capture the dewetting of aggregates of MDA-MB-231 human breast cancer cells [10], wettability and shape fluctuations of drug-resistant OVCAR-3 cancer cell aggregates [44], and the chemotaxis of swarms of *D. discoideum* [8]. We have thereby predicted how these biological processes emerge from active intermittent cell attachments.

Our model provides a coarse-grained description of the complex processes through which cells physically interact with their surrounding environment. The model is readily extendable to include further biological processes that have been found to impact self-organization, such as cell deformability [53, 54], cell growth and division [55] and contact inhibition of locomotion [26]. Broadly, our modeling framework can be applied to understand how cells dynamically regulate their physical interactions with their environment, to tune collective dynamics at the functionally relevant scale of multicellular assemblies.

## METHODS

### Model for intermittent attachments

The model for intermittent cell-cell and cell-surface attachments works as follows. Each cell can potentially form an attachment with any other cell that lies within an annular region, which is demarcated by an inner radius (*r*_1_) denoting the effective size of the cell, and an outer radius (*r*_2_) corresponding to the maximum length of a protrusion (Fig. 1C). To form an attachment, the cell randomly chooses another cell (or fixed surface sphere; see Fig. 1C,D) within this region; each cell within the attachment distance has an equal probability of being chosen (unless the direction of attachment is biased by chemotaxis; see Supplementary Materials). When a cell forms an attachment to another cell or surface sphere, it forms a spring which pulls the two cells towards each other with force given by Eq. (2) and Eq. (3) for cell-cell and cell-surface attachment forces, respectively. For cellsurface attachments, the surface sphere is not allowed to move. During the course of an attachment, if the distance between the two cells is less than or equal to *r*_1_, the attachment force becomes zero, and if the distance becomes greater than *r*_2_, the attachment is broken. Each spring-like force remains active for a duration of time before the attachment disappears and another is instantaneously formed using the same algorithm; therefore cell motility is persistent only during the lifetime of a single attachment. The duration of each attachment is sampled from a Gaussian distribution centered around *T*_*a*_, with variance *T*_*v*_, which is introduced to account for the variability in the timescale for cell-cell and cell-surface attachments that is observed in single-cell experiments [56].

All cells can form only one attachment or protrusion at any given time, but any cell can exchange forces with multiple other cells due to its own protrusion and those of other cells. Computationally, at any given point in time, we define the set of interaction partners (*P*_*i*_ in Eq. (2)) of each cell as all the cells that are exchanging mutually attractive forces with that cell. Our model is inherently non-equilibrium, because energy is injected each time a new attachment forms; when new springs form, they are extended and therefore possess elastic energy. In biological terms, this energy generation reflects force generation by the actin cytoskeleton through cellular metabolism. Note that the equations of motion of cells in our model, given by Eq. (4), do not include any noise term, which means that any kind of diffusion seen in the simulations emerges from the intermittent cell-cell attachments. Stochasticity in the model is caused by the random choice of other cells when a cell forms a new attachment.

### Parameter estimation

We estimate both the attachment force *K*_*a*_ and average attachment duration *T*_*a*_ from single-cell experimental data. Specifically, we estimate *K*_*a*_ from previous traction force microscopy measurements [57], and *T*_*a*_ from previous experiments analyzing the trajectories of individual cells [58] and imaging the dynamics of individual protrusions [56]. We estimate the value of the mobility parameter *µ* in Eq. (4) by computing the ratio of cell speed and magnitude of traction force from single-cell experimental data. The values of these, and other model parameters, including explanations of how they are estimated, are given in Table S1 in the Supplementary Materials.

When simulating our cell-based model, we consider the model parameters to be identical across all cells within the population, i.e. the population of cells is assumed to be phenotypically homogeneous. Therefore, we do not consider examples such as leader-follower dynamics observed in wound healing [59], where the dynamics are shaped by phenotypic heterogeneity within the population. We also neglect the effects of cell growth and division, which means that the total number and volume occupied by the cells is conserved in all the simulations.

## DATA AND SOFTWARE AVAILABILITY

All data are contained within the manuscript. Source code and initial conditions to reproduce simulation results are freely available at https://github.com/dpp98/Interm-attach-cells-3D

## ACKNOWLEDGMENTS

The authors acknowledge useful discussions with Shyamili Goutham, Lucija Mijanovic, David Versluis, Daniel Watson, Jonathan Chubb, Benjamin Walker and Mohit Dalwadi. We also thank Nir Gov for helpful suggestions on the text. P.P. was supported by a UK Research and Innovation (UKRI) Future Leaders Fellowship (MR/V022385/1). R.H.I. was supported by grants from the Medical Research Council (MR/X000702/1) and Wellcome Trust (221786/Z/20/Z). This work was supported by the Royal Society International Exchanges award IES\R1\241024 (R.B. and P.P.).

For the purpose of open access, the authors have applied a creative commons attribution (CC BY) licence to any author accepted manuscript version arising.

## References

[1] K. J. Cheung and A. J. Ewald, A collective route to metastasis: Seeding by tumor cell clusters, Science 352, 167 (2016).

[2] P. Friedl and D. Gilmour, Collective cell migration in morphogenesis, regeneration and cancer, Nature Reviews Molecular Cell Biology 10, 445 (2009).

[3] Y. Xiao, R. Riahi, P. Torab, D. D. Zhang, and P. K. Wong, Collective cell migration in 3D epithelial wound healing, ACS nano 13, 1204 (2019).

[4] A. G. Fletcher, M. Osterfield, R. E. Baker, and S. Y. Shvartsman, Vertex models of epithelial morphogenesis, Biophysical Journal 106, 2291 (2014).

[5] E. Scarpa and R. Mayor, Collective cell migration in development, Journal of Cell Biology 212, 143 (2016).

[6] T. J. Bartosh, J. H. Ylöstalo, A. Mohammadipoor, N. Bazhanov, K. Coble, K. Claypool, R. H. Lee, H. Choi, and D. J. Prockop, Aggregation of human mesenchymal stromal cells (MSCs) into 3D spheroids enhances their antiinflammatory properties, Proceedings of the National Academy of Sciences 107, 13724 (2010).

[7] T. R. Huycke, T. J. Häkkinen, H. Miyazaki, V. Srivastava, E. Barruet, C. S. McGinnis, A. Kalantari, J. Cornwall-Scoones, D. Vaka, Q. Zhu, et al., Patterning and folding of intestinal villi by active mesenchymal dewetting, Cell 187, 3072 (2024).

[8] H. Z. Ford, G. L. Celora, E. R. Westbrook, M. P. Dalwadi, B. J. Walker, H. Baumann, C. J. Weijer, P. Pearce, and J. R. Chubb, Pattern formation along signaling gradients driven by active droplet behavior of cell swarms, Proceedings of the National Academy of Sciences 122, e2419152122 (2025).

[9] G. Malet-Engra, W. Yu, A. Oldani, J. Rey-Barroso, N. S. Gov, G. Scita, and L. Dupré, Collective cell motility promotes chemotactic prowess and resistance to chemorepulsion, Current Biology 25, 242 (2015).

[10] C. Pérez-González, R. Alert, C. Blanch-Mercader, M. Gómez-González, T. Kolodziej, E. Bazellieres, J. Casademunt, and X. Trepat, Active wetting of epithelial tissues, Nature Physics 15, 79 (2019).

[11] S. Douezan, K. Guevorkian, R. Naouar, S. Dufour, D. Cuvelier, and F. Brochard-Wyart, Spreading dynamics and wetting transition of cellular aggregates, Proceedings of the National Academy of Sciences 108, 7315 (2011).

[12] G. Beaune, T. V. Stirbat, N. Khalifat, O. Cochet-Escartin, S. Garcia, V. V. Gurchenkov, M. P. Murrell, S. Dufour, D. Cuvelier, and F. Brochard-Wyart, How cells flow in the spreading of cellular aggregates, Proceedings of the National Academy of Sciences 111, 8055 (2014).

[13] R. A. Foty and M. S. Steinberg, Differential adhesion in model systems, Wiley Interdisciplinary Reviews: Developmental Biology 2, 631 (2013).

[14] C. Blanch-Mercader, R. Vincent, E. Bazellières, X. Serra-Picamal, X. Trepat, and J. Casademunt, Effective viscosity and dynamics of spreading epithelia: a solvable model, Soft Matter 13, 1235 (2017).

[15] D. Bonazzi, V. L. Schiavo, S. Machata, I. Djafer-Cherif, P. Nivoit, V. Manriquez, H. Tanimoto, J. Husson, N. Henry, H. Chaté, et al., Intermittent pili-mediated forces fluidize Neisseria meningitidis aggregates promoting vascular colonization, Cell 174, 143 (2018).

[16] H. Alston, A. O. Parry, R. Voituriez, and T. Bertrand, Intermittent attractive interactions lead to microphase separation in nonmotile active matter, Physical Review E 106, 034603 (2022).

[17] S. Ramaswamy, Active fluids, Nature Reviews Physics 1, 640 (2019).

[18] D. Bi, X. Yang, M. C. Marchetti, and M. L. Manning, Motility-driven glass and jamming transitions in biological tissues, Physical Review X 6, 021011 (2016).

[19] J. Rozman, K. Chaithanya, J. M. Yeomans, and R. Sknepnek, Vertex model with internal dissipation enables sustained flows, Nature Communications 16, 530 (2025).

[20] S. C. Kammeraat, Y.-E. Keta, P. Appleton, I. P. Newton, T. B. Liverpool, R. Sknepnek, I. Näthke, and S. Henkes, Correlated cell movements drive epithelial finger formation (2025), arXiv:2508.01046.

[21] S. Kim, R. Amini, S.-T. Yen, P. Pospíšil, A. Boutillon, A. Deniz, and O. Campàs, A nuclear jamming transition in vertebrate organogenesis, Nature Materials 23, 1592 (2024).

[22] F. Beroz, J. Yan, Y. Meir, B. Sabass, H. A. Stone, B. L. Bassler, and N. S. Wingreen, Verticalization of bacterial biofilms, Nature Physics 14, 954 (2018).

[23] R. Hartmann, P. K. Singh, P. Pearce, R. Mok, B. Song, F. Díaz-Pascual, J. Dunkel, and K. Drescher, Emergence of three-dimensional order and structure in growing biofilms, Nature Physics 15, 251 (2019).

[24] D. Oriola, M. Marin-Riera, K. Anlaş, N. Gritti, M. Sanaki-Matsumiya, G. Aalderink, M. Ebisuya, J. Sharpe, and V. Trivedi, Arrested coalescence of multi-cellular aggregates, Soft Matter 18, 3771 (2022).

[25] E. Palsson and H. G. Othmer, A model for individual and collective cell movement in *Dictyostelium* discoideum, Proceedings of the National Academy of Sciences 97, 10448 (2000).

[26] B. Smeets, R. Alert, J. Pešek, I. Pagonabarraga, H. Ramon, and R. Vincent, Emergent structures and dynamics of cell colonies by contact inhibition of locomotion, Proceedings of the National Academy of Sciences 113, 14621 (2016).

[27] T. D. Pollard and G. G. Borisy, Cellular motility driven by assembly and disassembly of actin filaments, Cell 112, 453 (2003).

[28] A. J. Ridley, M. A. Schwartz, K. Burridge, R. A. Firtel, M. H. Ginsberg, G. Borisy, J. T. Parsons, and A. R. Horwitz, Cell migration: integrating signals from front to back, Science 302, 1704 (2003).

[29] T. Lämmermann, B. L. Bader, S. J. Monkley, T. Worbs, R. Wedlich-Söldner, K. Hirsch, M. Keller, R. Förster, D. R. Critchley, R. Fässler, et al., Rapid leukocyte migration by integrin-independent flowing and squeezing, Nature 453, 51 (2008).

[30] W.-h. Guo and Y.-l. Wang, A three-component mechanism for fibroblast migration with a contractile cell body that couples a myosin II–independent propulsive anterior to a myosin II–dependent resistive tail, Molecular Biology of the Cell 23, 1657 (2012).

[31] H. Yamaguchi and J. Condeelis, Regulation of the actin cytoskeleton in cancer cell migration and invasion, Biochimica et Biophysica Acta (BBA)-Molecular Cell Research 1773, 642 (2007).

[32] R. J. Petrie and K. M. Yamada, Multiple mechanisms of 3D migration: the origins of plasticity, Current Opinion in Cell Biology 42, 7 (2016).

[33] C. E. Chan and D. J. Odde, Traction dynamics of filopodia on compliant substrates, Science 322, 1687 (2008).

[34] A. J. Ridley, Rho GTPase signalling in cell migration, Current Opinion in Cell Biology 36, 103 (2015).

[35] S. Sart, A.-C. Tsai, Y. Li, and T. Ma, Three-dimensional aggregates of mesenchymal stem cells: cellular mechanisms, biological properties, and applications, Tissue Engineering Part B: Reviews 20, 365 (2014).

[36] G. Giannone, B. J. Dubin-Thaler, O. Rossier, Y. Cai, O. Chaga, G. Jiang, W. Beaver, H.-G. Döbereiner, Y. Freund, G. Borisy, et al., Lamellipodial actin mechanically links myosin activity with adhesion-site formation, Cell 128, 561 (2007).

[37] A. P. Thompson, H. M. Aktulga, R. Berger, D. S. Bolintineanu, W. M. Brown, P. S. Crozier, P. J. i. t. Veld, A. Kohlmeyer, S. G. Moore, T. D. Nguyen, R. Shan, M. J. Stevens, J. Tranchida, C. Trott, and S. J. Plimpton, LAMMPS - a flexible simulation tool for particle-based materials modeling at the atomic, meso, and continuum scales, Computer Physics Communications 271, 108171 (2022).

[38] H. Z. Ford, A. Manhart, and J. R. Chubb, Controlling periodic long-range signalling to drive a morphogenetic transition, Elife 12, e83796 (2023).

[39] R. A. Foty and M. S. Steinberg, Surface tensions of embryonic tissues predict their mutual envelopment behavior, Development 120, 853 (1994).

[40] J. G. Méndez-Bermúdez, I. Guillén-Escamilla, G. A. Méndez-Maldonado, and J. A. D. Alva-Tamayo, Argon force field revisited: a molecular dynamic study, Journal of Physics Communications 6, 041002 (2022).

[41] M. P. Allen and D. J. Tildesley, Computer Simulation of Liquids (Oxford University Press, 2017).

[42] G. Beaune, C. Blanch-Mercader, S. Douezan, J. Dumond, D. Gonzalez-Rodriguez, D. Cuvelier, T. Ondarçuhu, P. Sens, S. Dufour, M. P. Murrell, et al., Spontaneous migration of cellular aggregates from giant keratocytes to running spheroids, Proceedings of the National Academy of Sciences 115, 12926 (2018).

[43] A. Ravasio, A. P. Le, T. B. Saw, V. Tarle, H. T. Ong, C. Bertocchi, R.-M. Mège, C. T. Lim, N. S. Gov, and B. Ladoux, Regulation of epithelial cell organization by tuning cell–substrate adhesion, Integrative Biology 7, 1228 (2015).

[44] S. Goutham, A. Gopikrishnan, B. Harshavardhan, N. Venas, R. Sabarinathan, M. K. Jolly, and R. Bhat, Collective amoeboid dynamics drives colonization of drug-resistant ovarian cancer cells, bioRxiv, 2024 (2024).

[45] A. J. Muinonen-Martin, O. Susanto, Q. Zhang, E. Smethurst, W. J. Faller, D. M. Veltman, G. Kalna, C. Lindsay, D. C. Bennett, O. J. Sansom, et al., Melanoma cells break down LPA to establish local gradients that drive chemotactic dispersal, PLOS Biology 12, e1001966 (2014).

[46] J. Stock and A. Pauli, Self-organized cell migration across scales–from single cell movement to tissue formation, Development 148 (2021).

[47] L. Tweedy, D. A. Knecht, G. M. Mackay, and R. H. Insall, Self-generated chemoattractant gradients: attractant depletion extends the range and robustness of chemotaxis, PLOS Biology 14, e1002404 (2016).

[48] E. F. Keller and L. A. Segel, Traveling bands of chemotactic bacteria: a theoretical analysis, Journal of theoretical biology 30, 235 (1971).

[49] M. Sanoria, G. Malet-Engra, G. Scita, N. Gov, and A. Gopinathan, Chemotaxis-driven instabilities govern size, shape and migration efficiency of multicellular clusters, arXiv preprint arXiv:2506.00310 (2025).

[50] A. Mongera, P. Rowghanian, H. J. Gustafson, E. Shelton, D. A. Kealhofer, E. K. Carn, F. Serwane, A. A. Lucio, J. Giammona, and O. Campàs, A fluid-to-solid jamming transition underlies vertebrate body axis elongation, Nature 561, 401 (2018).

[51] R. J. Tetley, M. F. Staddon, D. Heller, A. Hoppe, S. Banerjee, and Y. Mao, Tissue fluidity promotes epithelial wound healing, Nature Physics 15, 1195 (2019).

[52] P. Schaap, Evolutionary crossroads in developmental biology: *Dictyostelium discoideum*, Development 138, 387 (2011).

[53] M. R. Nejad and J. M. Yeomans, Coarse-graining dense, deformable active particles (2025), arXiv:2501.07280.

[54] A. Torres-Sánchez, M. Kerr Winter, and G. Salbreux, Interacting active surfaces: A model for three-dimensional cell aggregates, PLOS Computational Biology 18, e1010762 (2022).

[55] O. Hallatschek, S. S. Datta, K. Drescher, J. Dunkel, J. Elgeti, B. Waclaw, and N. S. Wingreen, Proliferating active matter, Nature Reviews Physics 5, 407 (2023).

[56] N. Andrew and R. H. Insall, Chemotaxis in shallow gradients is mediated independently of PtdIns 3-kinase by biased choices between random protrusions, Nature Cell Biology 9, 193 (2007).

[57] K. Inouye and I. Takeuchi, Motive force of the migrating pseudoplasmodium of the cellular slime mould *Dictyostelium* discoideum, Journal of Cell Science 41, 53 (1980).

[58] L. Bosgraaf and P. J. Van Haastert, The ordered extension of pseudopodia by amoeboid cells in the absence of external cues, PLOS one 4, e5253 (2009).

[59] M. Vishwakarma, B. Thurakkal, J. P. Spatz, and T. Das, Dynamic heterogeneity influences the leader–follower dynamics during epithelial wound closure, Philosophical Transactions of the Royal Society B 375, 20190391 (2020).

