## Supplementary Text for "Intermittent attachments form three-dimensional cell aggregates with emergent fluid properties"

This PDF file contains Figures S1-S5, Supporting Text, Table S1 summarizing the simulation parameters, and Legends for Movies S1-S4

### Additional Figures

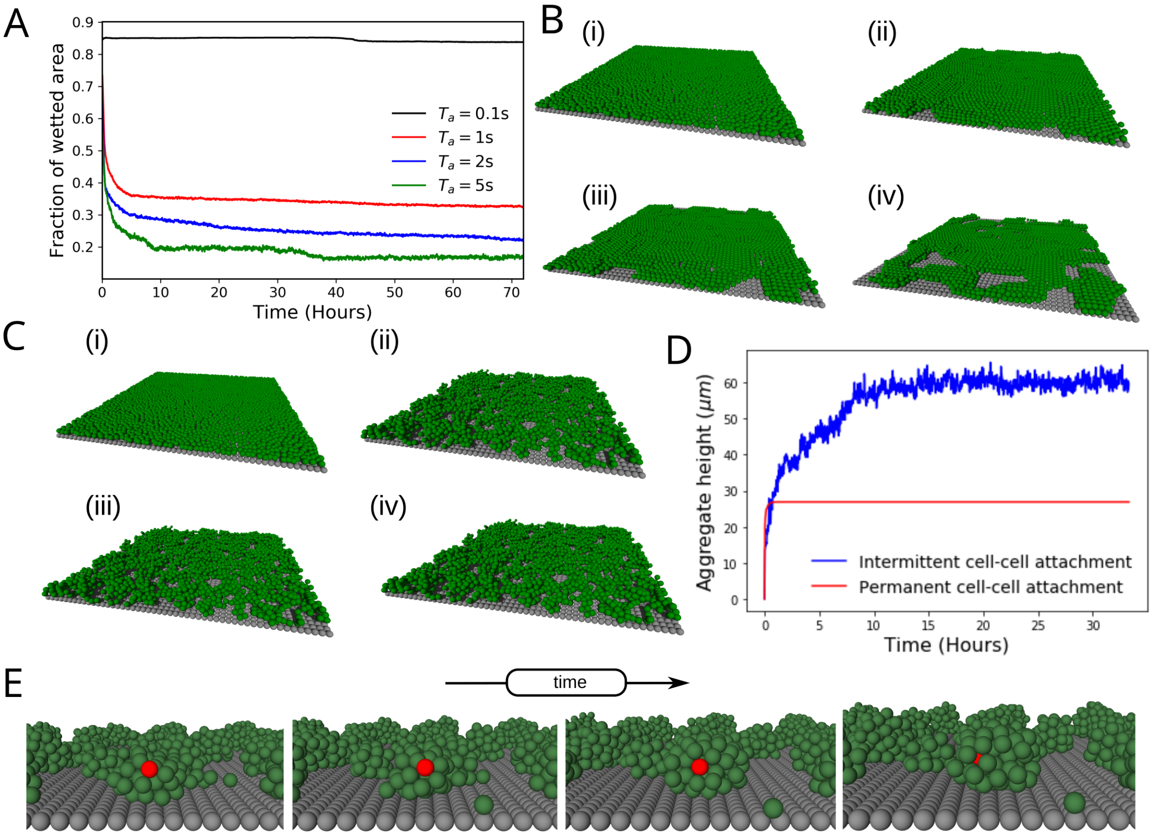


Figure S1: (A) Time-evolution of the fraction of the surface area wetted by the cells for an initial monolayer of cells on a surface. Different colors correspond to simulations with distinct values of attachment time $T_a$. (B) Simulation snapshots showing a monolayer of cells that has a very short attachment time ($T_a = 0.1s$), corresponding to the black line in A. Time increases from (i) to (iv). (C) Snapshots of a simulation showing that a monolayer of cells having permanent cell-cell attachments is unable to form droplet-like aggregates seen in Fig. 2A of the main text. The average cell-surface attachments last for the default value of 10s. (D) Variation in height of the aggregate with time for intermittent and permanent cell-cell attachments, showing a comparison of the formation of 3D surface-associated aggregates. For both cases, the cell-surface attachments last for an average of 10s. The initial increase in height observed for permanent cell–cell attachment arises because, at the start of the simulation, cells are assumed to be unattached. (E) Single-cell tracking of cells within a aggregate shows that a cell can rearrange its position within the aggregate. Here the value of $w_f$ was taken to be equal to 0.25, and the monolayer packing fraction in the initial condition was equal to 0.5.

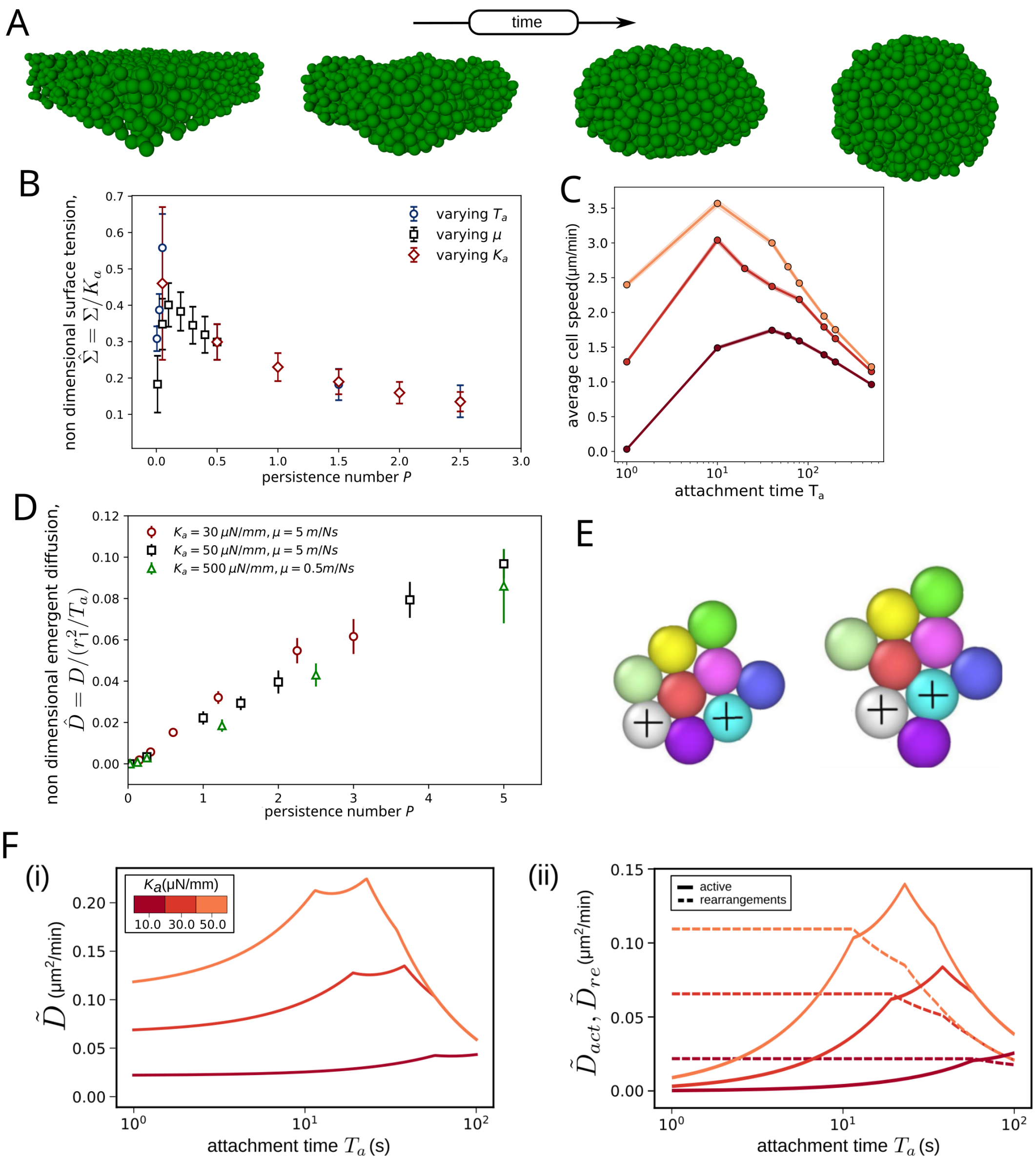


Figure S2: (A) Simulation snapshots showing shape relaxation of an aggregate which was initially shaped as a cuboid and spontaneously minimizes its surface area to form a sphere. (B) Variation of non dimensional surface tension, measured using the parallel plate compression test (Fig. 3A) with the dimensionless persistence number. (C) Variation of ensemble averaged cell speed with attachment time. (D) Variation of non dimensional emergent diffusivity with persistence number. The results shown in C and D are obtained from simulations described in Fig. 3D.(E) Simulation snapshots showing rearrangements within a 2D cell aggregate. Here, only the gray and light-blue colored cells are active partners and experience an active force (see Eq. (2) in the main text); the violet and red colored cells instead move passively due to volume exclusion. This example shows how active cell-cell attachments can lead to both active movement and passive rearrangements within a cell aggregate. (F) (i) Emergent diffusion coefficient $\tilde{D}$ as a function of the attachment time $T_a$ as predicted from the reduced 1D model, when the maximum permissible length of attachment $r_2$ equals $60\mu m$. Different colors distinguish predictions for different values of the attachment strength $K_a$. (ii) Contributions from the active attachments $\tilde{D}_{act}$ and rearrangements $\tilde{D}_{re}$ induced by the active attachments to the emergent diffusion, respectively.

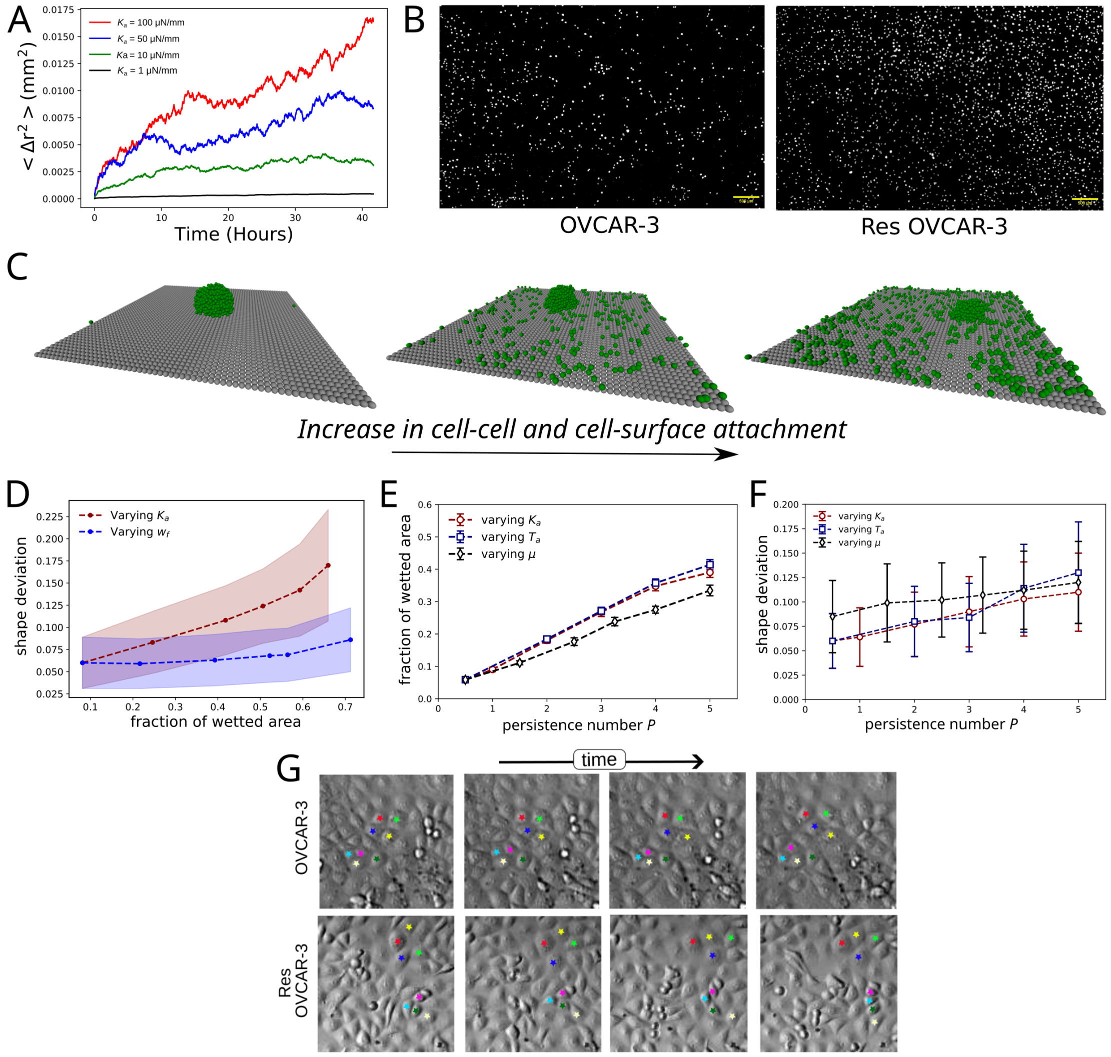


Figure S3: (A) Mean Squared displacement vs. time for single cells, for different values of $K_a$ showing that an increase in $K_a$ increases the motility of single cells. (B) Trans-well migration assay for drug-resistant ovarian cancer cells, obtained from Ref. 44, (licensed under CC BY-NC-ND 4.0). This is a well-established technique in which single-cells' motility is studied by measuring the ability of cells to cross a porous medium. The panels illustrate the difference in the number of drug-sensitive (left) and drug-resistant (right) cells that have been able to cross the porous medium during the time of the experiment. The larger number of cells in the right panels, therefore, indicates that the resistant OVCAR-3 cells are more motile as compared to the non-resistant ones. (C) 3D snapshots at steady state, illustrating the wetting of aggregates of cells with increasing attachment strength, $K_a$: (left) $K_a = 1$ dyne/mm (same as Fig 5D); (middle) $K_a = 10$ dyne/mm; (right) $K_a = 25$ dyne/mm corresponding to the simulations shown in Fig. 5C,D of the main text. (D) Plot of aggregates equilibrium shape deviation as a function of their equilibrium wetting. We compare results for two different drivers of wetting, *i.e.*, by increasing the strength of cell-surface relative to cell-cell attachments ($w_f$), and by increasing the strength of both cell-surface and cell-cell attachments ($K_a$). (E) Variation in the fraction of the surface area wetted at steady state with the persistence number $P$ when $w_f = 1$. (F) Variation in shape deviation with the persistence number Ps shows a monotonic increase in the magnitude of shape deviation with increase in $P$. (G) Time-lapse imaging of a monolayer of OVCAR-3 ovarian cancer cells (top), and their drug-resistant counterpart cells (Res OVCAR-3) (bottom), seeded on a glass substrate, where the relative position of two pairs of four neighboring cells is marked (using red, blue, green, and yellow asterisks) to enable the visualization of their motility and degree of inter-positional rearrangement, obtained from [44] (licensed under CC BY-NC-ND 4.0). Comparison of the top and bottom panels reveals that aggregates of drug-resistant ovarian cancer cells exhibit greater fluidity as compared to the sensitive cells, as they can be seen to exhibit greater cell-cell rearrangements.

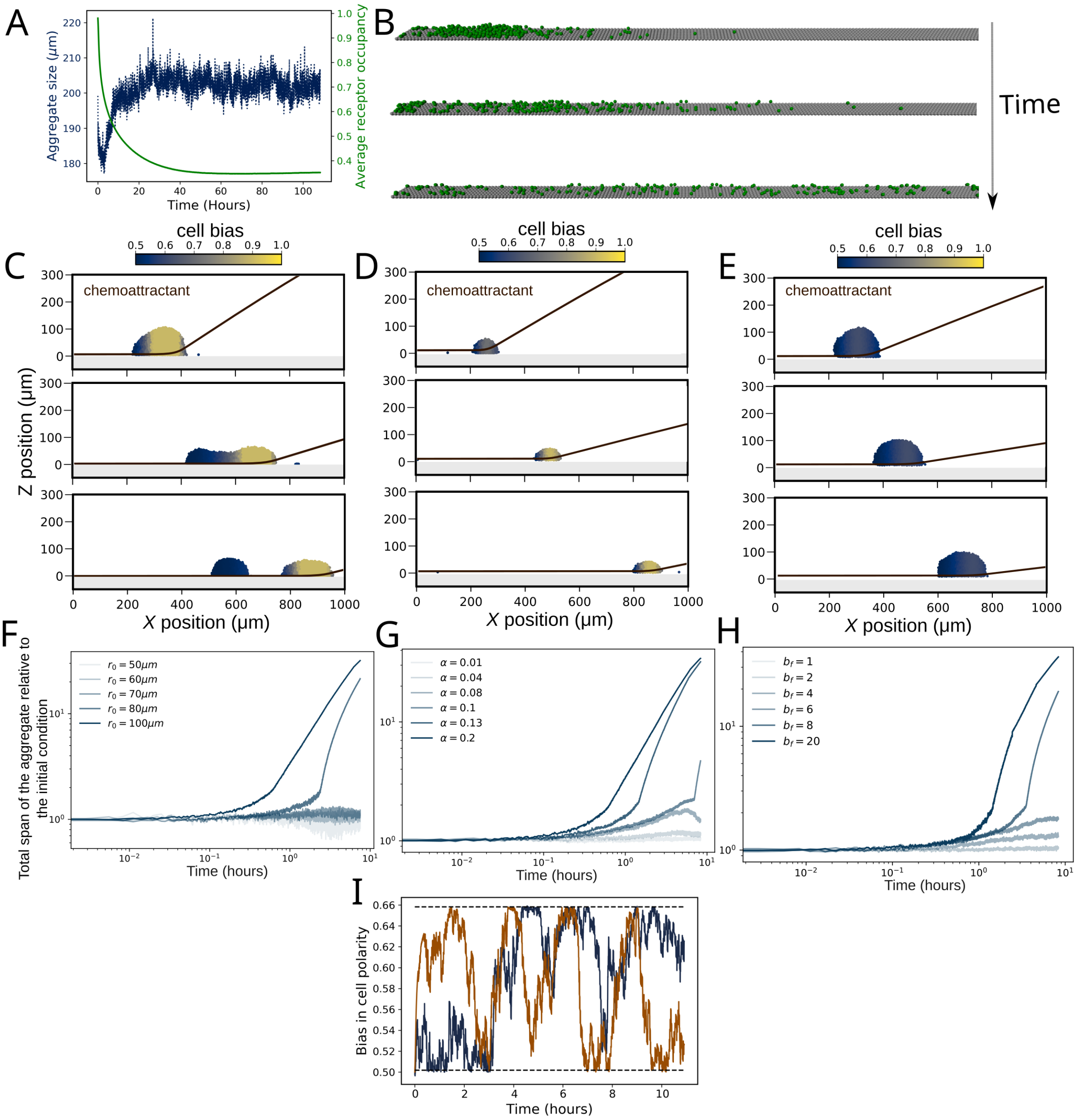


Figure S4: (A) Temporal variation in aggregate size and average cell receptor occupancy shows that the migrating aggregate eventually attains a traveling state with a stationary speed and shape. Note that fluctuation in the aggregate size is on the scale of a cell radius. (B) Snapshots of a simulation of self-generated chemotaxis of a dilute population of cells, where the cells are shown to migrate individually on the surface, as opposed to the collective aggregate migration shown in Fig. 6B in the main text. This results from the combined effect of higher adhesion to the surface ($w_f$) and higher chemotactic bias at the front part of the aggregate. (C,D) Effect of decreasing the initial aggregate size on splitting: (C) default value (1470 cells, corresponding to an initial aggregate radius $r_0 = 100\mu m$); (D) 367 cells corresponding to $r_0 = 50\mu m$. All other parameters are specified in Table S1. (C,E) Effect of decreasing the maximum rate of chemoattractant consumption ($\alpha$) on splitting: (C) $\alpha = 0.20$; (E) $\alpha = 0.01$, keeping all other parameters the same. (F-H) Variation of total aggregate span with time for different values of (F) initial aggregate size, (G) chemoattractant consumption rate, and (H) chemotactic sensitivity. (I) Time evolution of the bias experienced by two randomly chosen cells within a migrating aggregate (same simulations as in Fig. 6B). The black dotted lines denote the minimum and maximum values of bias within the aggregate. The blue and brown colored curves suggest that cells can experience the entire range of bias available within the aggregate.

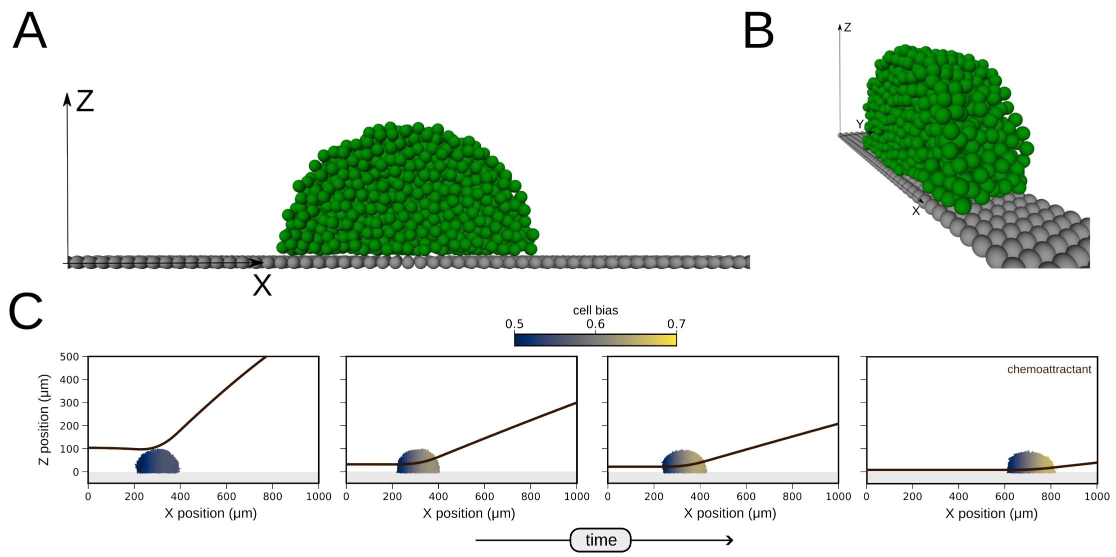


Figure S5: A top side view and perspective view of the self-generated chemotaxis simulation setup are shown in A and B, respectively. (C) Snapshots of a simulation showing the generation of a gradient in chemoattractant concentration, and collective migration of the cell aggregate due to self-generated chemotaxis (same as Fig. 6B).

### Supporting Text

#### S1 Cell-based simulations.

##### S1.1 Non-dimensionalization of active intermittent attachments model

We set the length and time scales equal to $r_1$ and $T_a$ respectively, and use the product of the attachment strength $K_a$ and $r_1$ as the force scale. This gives us the following non-dimensionalized variables for distance, time, and force, respectively: $\tilde{x} = \frac{x}{r_1}$, $\tilde{t} = \frac{t}{T_a}$, $\tilde{F} = \frac{F}{K_a r_1}$. Substituting the dimensionalized variables into equation (4) of the main text, we get the following non-dimensional equation for the motion of the cells,

$$\frac{d\tilde{x}_i}{d\tilde{t}} = \underbrace{\mu K_a T_a}_{P} \hat{F}_i, \tag{1}$$

where $\tilde{F}_i = \tilde{F}_i^{WCA} + \tilde{F}_i^{A} + \tilde{F}_i^{S}$. The non-dimensional forces are

$$\tilde{F}_i^{WCA} = \left[2\left(\frac{\sigma}{r_1\tilde{r}_{ij}}\right)^{12} - \left(\frac{\sigma}{r_1\tilde{r}_{ij}}\right)^{6}\right]\frac{24\varepsilon\hat{\tilde{r}}_{ij}}{K_a r_1^2 \tilde{r}_{ij}}, \ \tilde{r}_{ij} \leq 2^{\frac{1}{6}}(\sigma/r_1) \tag{2}$$

$$\tilde{F}_i^{A} = \sum_{j\in P_i}(1-\tilde{r}_{ij})\hat{\tilde{r}}_{ij}, \quad \tilde{r}_{ij} \geq 1 \tag{3}$$

$$\tilde{F}_i^{S} = \sum_{j\in S_i} w_f(1-\tilde{r}_{ij})\hat{\tilde{r}}_{ij}, \quad \tilde{r}_{ij} \geq 1. \tag{4}$$

The non-dimensional numbers in the problem are as follows. The key non-dimensional parameter that we identify is $P = \mu K_a T_a$, which we call the persistence number; this approximately quantifies the distance moved by a cell per unit time owing to active forces, along the direction the attachment (until the spring reaches its natural length). It emerges naturally from this non-dimensionalization and plays a key role in governing the emergent dynamics of cell aggregates (as shown in Figs S2B,D, and S3E-F). The other non–dimensional parameters are $\epsilon/(K_a r_1^2)$, which quantifies the strength of volume exclusion, $\sigma/r_1$, which quantifies the range of volume exclusion, $w_f$, which quantifies the relative strength of cell-surface and cell-cell attachments, and $r_2/r_1$, which quantifies the maximum range of active attachments. The only parameters modified in this study are $\mu$, $K_a$, $T_a$, and $w_f$; the other parameters are fixed at biologically relevant values (see Table S1)

Based on these scalings, the emergent surface tension is non-dimensionalized using the attachment strength, leading to $\hat{\Sigma} = \frac{\Sigma}{K_a}$ and the non-dimensionalized emergent diffusion coefficient is given by $\hat{D} = \frac{D}{(r_1^2/T_a)}$.

##### S1.2 Initialization

We here complement the Method section in the main text with additional details on how we initialize model simulations.

The initial configuration of cells was chosen according to the physical phenomena under investigation. For Figs. 2A-C and S1A-E, we started from a monolayer of cells similar to the left-most panel of Fig. 2A. For estimating the surface tension in Figs. 3A-C and S2B, we started from a spherically shaped aggregate that was slowly compressed by moving the two plates closer to each other (see schematic in Fig. 3A). The aggregate was allowed to relax to a steady state after which the relevant quantities were calculated (see Eq. (5) of the main text). For Figs. 3D-G and S2C-D, we used a simulated a cuboid with periodic boundary conditions, as depicted in Fig. 3D. The initial condition for Fig. 4A was generated by first letting a hemispherical shaped aggregate spread over a surface by taking a value of $w_f = 3.5$. The steady-state configuration from this simulation was taken as the initial condition for Figs. 4A and 4B. For Fig. S3A, we started with 9 cells placed on a $3 \times 3$ lattice that has a separation of 200 $\mu m$ between consecutive cells to ensure that cell-cell interactions do not influence the single cell motility considered in this figure. In Figs. 4D, 5C-E, and S3C-F, we started with a hemispherical shaped aggregate over a surface, and, subsequently, let it evolve with time. For Figs. 6B-E and S4A-I, we started from a semi-cylindrically shaped aggregate, which satisfied the periodic boundary conditions along the $y$-direction. To eliminate any stochastic artefact, all the quantitative results presented in the paper were obtained averaging over a large ensemble of steady states consisting of at least 1000 configurations that are well separated in time, and also validated our results with multiple independent simulation runs.

Since the model includes a WCA repulsion potential, it is important to minimize the overlap in the initial position of the cells to avoid instabilities due to unrealistically high values of the repulsive forces (see Eq. (1) in the main text). First, we specify the geometry of the bounding box for a given initial condition. We then calculate the number of cells that are required in the given volume for the desired cell density, and use rejection sampling to choose the cell positions and reject cells that have a significant overlap. For example, for the initial condition corresponding to a cell monolayer, we randomly choose the Cartesian coordinates of the cell centroids within the 2D rectangular domain given by the $x$ and $y$ limits of the boundary, and reject cells that overlap with previously chosen cells. This process is iterated until we have the desired number of cells in the given volume. Similarly, for the hemispherical and semi-cylindrical shaped initial conditions, we use spherical and cylindrical polar coordinates, respectively, for specifying the domain boundary and sampling the positions of the cell centroids within it.

#### S1.3 Measuring shape deviation of aggregates

We here describe how we estimate the deviation from circularity of simulated aggregates,*i.e.*, shape deviation, reported in Figs. 2E, S2D and S2F. We fit a convex hull to the cluster of cells, see Fig. 2D in the main text for an example. We measure the centroid of the cluster and compute the distance of each point in the convex hull from the centroid. We measure the standard deviation of these distances and normalize it with the average distance of the convex hull from the centroid. We take an average of this deviation across all time steps after the aggregate has reached steady state, and also report the variation of the quantity over time (denoted by the error bar in Fig. 2E).

#### S1.4 Modeling self-generated chemotaxis

We summarize here the mathematical and implementation details of the simulations for studying collective chemotaxis of cell aggregates, due to self-generated chemotaxis.

We consider a cell aggregate on a surface as shown in Fig. S5, with periodic boundary conditions along the $y$-direction, and an unconstrained boundary along the $z$-direction. We assume a 1D chemoattractant concentration profile $C = C(x,t)$ which evolves according to the following equation:

$$\frac{\partial C}{\partial t} = D_c \nabla^2 C - \sum_{i=1}^{N} \alpha \frac{C(X_i,t)}{C(X_i,t) + k_M} \delta(x, X_i), \tag{5}$$

where $D_c$ denotes the diffusivity of the chemoattractant, $N$ denotes the number of cells in the simulation, $X_i$ denotes the $x$-coordinate of the $i^{\text{th}}$ cell at time $t$, $\alpha$ denotes the maximum rate of degradation of the chemoattractant by a cell, $k_M$ denotes the Michaelis-Menten constant, and $\delta$ denotes the approximate dirac-delta function. Here we assume that the chemoattractant is degraded by the cells locally, as represented by the second term on the RHS of Eq. (5). We assume an initially uniform concentration profile such that $C(x,0) = C_0, \quad \forall x \in [0,L]$, along with a no-flux boundary condition at $x = 0$, and a Dirichlet boundary condition at $x = L$, such that $C(L,t) = C_0$ for all $t > 0$. We discretise Eq. (5) on a set of uniformly spaced points, $x_j = j\Delta x$ with $\Delta x = 10\mu$m, using central difference and advance it in time using the forward Euler scheme with a fixed time-step $\Delta t = 0.01$ s. The consumption term in the RHS of (5) is approximated by considering the decay for each cell to take place only at the two grid points that are adjacent to the position of the cell centroid, and the distance between two grid points is chosen to be consistent with the cell size (see Table S1).

A combination of diffusion and localized degradation of chemoattractant by the cells leads to the formation of a gradient in chemoattractant concentration, as shown in Fig. S5C. We assume cells respond to the gradient by biasing their polarity along the $x$ direction. If the position of a cell centroid lies between the grid points $x_j$ and $x_{j+1}$, i.e., $X_i \in (x_i, x_{j+1})$, then the bias experienced by the cells is given by

$$P_R = 0.5 + b_f \left[ \frac{C(x_{j+1})}{C(x_{j+1}) + K_d} - \frac{C(x_j)}{C(x_j) + K_d} \right], \tag{6}$$

where $b_f$ denotes the chemotactic sensitivity of the cells, and $K_d$ denotes the dissociation constant for the binding of the chemoattractant with receptors on the cell membrane. We constrain $P_R$ to be bounded between 0 and 1, such that it denotes the likelihood for a cell to form an attachment along the positive $x$ direction. When $P_R$ is equal to 0.5, the cell is equally likely to form an attachment both along positive and negative $x$. When $P_R$ is greater than 0.5, the cell is more likely to attach to another cell or a surface sphere that is located along the $+\hat{\boldsymbol{x}}$ direction with respect to its position. Similarly, $P_R < 0.5$ corresponds to an increase in the likelihood of forming an attachment along the $-\hat{\boldsymbol{x}}$ direction. The polarization of the cells along the $y$ and $z$ directions is assumed to be unaffected by the chemoattractant. This model of chemotaxis is based on a reduced one-dimensional model for self-generated chemotaxis that was introduced in [47].

### S2 1-D Model for emergent fluidity

We here outline the details of the reduced 1D model of cell movement via active intermittent attachments in dense cell populations used to derive Eqs. (8-9) in the main text and generate Figs. 4J and S2F.

**Set-up.** We model $n$ cells that form a 1D chain with periodic boundary conditions. Each cell touches its right and left neighbours, hence neighbouring cell centers are a distance of $\Delta x := 2r_1$ apart, where $r_1$ is the radius of one cell. Cell movement can happen either actively, where two interaction partners form a transient connection and pull on each other, or due to volume exclusion effects, where cells are moved out of the way by actively moving neighboring cells.

**Active movement.** A cell chooses its interaction partner from the $2n_I \in \mathbb{N}$ cells to its right or left, where $n_I = \lceil \frac{r_2 - r_1}{2r_1} \rceil$ is its reach in number of cells in each direction. Hence, each cell has $2n_I$ possible interaction partners, chosen with equal probability. Once established, the interaction lasts for an average of $T_a$ time units. Connected cells move in steps of $\Delta x$ towards each other until they 1) touch or 2) stop interacting, whichever happens earlier. If there is only one cell between them, one of the two partners is chosen at random to move. Given the mobility $\mu$ and attraction strength $K_a$ (see main text), the time it takes to move one cell diameter can be estimated by $T_d = \frac{2r_1}{\mu K_a\left(\frac{r1+r2}{2}\right)}$. This leads to an upper bound of steps a connected cell could move while interacting of $s := T_a/T_d$. However, $s$ will generally not be an integer. Therefore, we assume this upper bound to be $\lfloor s \rfloor$ with probability $q = 1 - (s - \lfloor s \rfloor)$ and $\lfloor s \rfloor + 1$ with probability $1 - q$. If interaction partners are $j \in \{1, \ldots, n_I\}$ cell diameters ($\Delta x$) away from each other, they would touch after $\frac{j-1}{2}$ steps, if they stay connected. Putting this together means connected cells move an average of

$$d_j := q \min \left\{ \frac{j-1}{2}, \lfloor s \rfloor \right\} + (1-q) \min \left\{ \frac{j-1}{2}, \lfloor s \rfloor + 1 \right\} \tag{7}$$

steps of size $\Delta x$.

**Cell Rearrangements** If an actively moving cell moves $\Delta x$ to the right(/left), it switches places with its right(/left) neighbour. These neighbours are therefore moved passively of a distance $\Delta x$ to the left(/right).

**Emergent diffusion.** We denote by $X_i$ the random variable describing the displacement of the $i$-th cell. For given $j \in \{1, \ldots, n_I\}$ the probability to be the left(/right) active partner is $p = \frac{2}{n}\frac{1}{2} = \frac{1}{n}$, which will move $X_i = \pm d_j \Delta x$ to the right(/left). The probability to be a passively moving left(/right) cell is the probability *not* to be an actively moving cell $1 - \frac{2}{n}$ times the probability to be in the path of an actively moving right(/left) cell $\frac{d_j}{n-2}$, i.e. $p = (1-\frac{2}{n})\frac{d_j}{n-2} = \frac{d_j}{n}$, which leads to movement of $\pm\Delta x$ during the course of the interaction of the actively moving cells. Together, this yields the expected displacement and variance of displacement:

$$\mathrm{E}[X_i] = \frac{n}{n_I}\sum_{j=1}^{n_I}\left(\frac{1}{n}(-d_j\Delta x) + \frac{1}{n}(d_j\Delta x) + \frac{d_j}{n}(-\Delta x) + \frac{d_j}{n}(\Delta x)\right) = 0,$$
$$\mathrm{E}[X_i^2] = \frac{n}{n_I}\sum_{j=1}^{n_I}\left(\frac{1}{n}(d_j\Delta x)^2 + \frac{1}{n}(d_j\Delta x)^2 + \frac{d_j}{n}(\Delta x)^2 + \frac{d_j}{n}(\Delta x)^2\right) = \frac{2\Delta x^2}{n_I}\sum_{j=1}^{n_I} d_j^2 + d_j.$$

This shows that the process indeed leads to emergent diffusion. Given that we described the dynamics over the course of one interaction lasting an average time of of $T_a$, we obtain for the mean square displacement until time $t$

$$\mathrm{MSD} = \frac{t}{T_a}\mathrm{E}[X_i^2] = 2Dt,$$

which finally leads to a diffusion constant of

$$D = \frac{(2r_1)^2}{T_a}\frac{1}{n_I}\sum_{j=1}^{n_I} d_j^2 + d_j, \tag{8}$$

where $d_j$ is defined by Eq. (7), $q = 1-(s-\lfloor s\rfloor)$, $s = \lfloor\frac{T_a}{T_d}\rfloor$, $n_I = \lceil\frac{r_2-r_1}{2r_1}\rceil$ and $T_d = \frac{2r_1}{\mu K_a\left(\frac{r1+r2}{2}\right)}$.

**Interpretation.** Formula (8) shows that the active contributions enter quadratically in $d_j$ and the contributions from rearrangements enter linearly in $d_j$, demonstrating their differential effect on the emergent diffusion. Further, we see that the attraction strength $K_a$ increases diffusion. The effect of the interaction time $T_a$ is more complicated, and we have two regimes

1. Long interaction times: $\lfloor\frac{T_a}{T_d}\rfloor > \frac{n_I-1}{2}$. The interaction time is not limiting $d_j$, and we obtain

   $$D = \frac{(r_1)^2}{T_a}\frac{2n_I^2+3n_I-5}{6}.$$

   In this case, longer interaction times do not lead to more movement during an individual interaction but rather decrease the number of different interactions up to a given time $t$. Hence, longer interaction times decrease the diffusion rate.

2. Short interaction times: $\lfloor\frac{T_a}{T_d}\rfloor < \frac{n_I-1}{2}$. Here the interaction time is limiting $d_j$, and longer interaction times lead to more movement during an individual's interactions.

### Simulation Parameters

| Parameter | Default Value(s) | Unit | Description |
|---|---|---|---|
| $K_a$ | 10.0 | $\mu N\ \cdot$mm$^{-1}$ | Strength of active force exerted by cells using pseudopodia [57]. |
| $\epsilon$ | $10^{-11}$ | Joule | Strength of the WCA repulsion potential, which is responsible for ensuring volume exclusion. The value of this parameter is chosen based on numerical stability and the accuracy of implementing volume exclusion. |
| $\sigma$ | $10.0/2^{1/6}$ | $\mu$m | Length scale of WCA repulsion, assuming cell size to be 10 $\mu$m. |
| $T_a$ | 10.0 | $s$ | Average time duration for which an attachment remains active. Estimated from single cell experiments [58]. |
| $T_v$ | $\frac{T_a}{10}$ | $s^2$ | Variance in the attachment time. |
| $\mu$ | 50 | $m \cdot N^{-1} \cdot s^{-1}$ | Mobility of the particle within the medium. Indirectly estimated from pseudopod traction and cell velocity, with the assumption of inertia-less motion. |
| $r_1$ | 10 | $\mu$m | Minimum distance between two particles for attraction. |
| $r_2$ | 30 | $\mu$m | Maximum distance between two particles for attraction. The values of $r_1$ and $r_2$ are estimated by considering the aspect ratio of single migrating cells from experiments [56]. |
| $w_f$ | 1.0 | non-dimensional | Relative force of cell-surface and cell-cell attachment. |
| $D_c$ | 200 | $\mu$m$^2\ \cdot s^{-1}$ | Diffusivity of chemoattractant, taken from [47]. |
| $k_M$ | 700 | nM | Michaelis-Menten constant for chemoattractant consumption, obtained from [47]). |
| $K_d$ | 20 | nM | Dissociation constant for binding of chemoattractant with the receptors on the cell surface, obtained from [47]. |
| $b_f$ | 10 | non-dimensional | Chemotactic sensitivity parameter (Assumed). |
| $L$ | 10 | mm | Length of the simulation domain for the chemotaxis simulations, chosen to be much larger than the aggregate size, to observe migration over a long distance. |
| $\alpha$ | 0.01 | nM $\cdot$s$^{-1}$ | Maximum rate of degradation of the chemoattractant (estimated from the rate of degradation of folate by *D. discoideum* [47]). |
| $\Delta x$ | 10.0 | $\mu$m | Grid spacing used to solve for the chemoattractant field chosen to be equal to the cell size for simplicity. |
| $C_0$ | 10 | $\mu$M | Initial value of the chemoattractant concentration field, in accordance with realistic experimental concentrations [47]. |
| $\Delta t$ | 0.01 | $s$ | Timestep for the simulation, chosen to ensure the invariance of the results to further decrease in timestep. |

Table S1: List of parameters used in the cell-based simulations along with their default value.

### Movies

#### Movie S1. Visualization of active intermittent attachments

Here we show an example of a small-sized aggregate with only 21 cells and 49 surface spheres. An attachment between two cells is colored in red and cell-surface attachments are colored blue.

#### Movie S2. Dewetting of a cell monolayer

Here we show an animation for the simulation corresponding to Fig. 2A where we can see that a monolayer of cells can spontaneously dewet a surface, leading to the formation of three-dimensional surface associated cell-aggregates.

#### Movie S3. Dewetting of a cell monolayer for different attachment times

Here we compare the dewetting of a monolayer of cells on a substrate, for four different values of attachment time, corresponding to Figure 3H.

#### Movie S4. Shape fluctuations of surface associated cell aggregates

Here we show two surface-associated cell aggregates at steady state for two different values of attachment strength (corresponding to Fig. 5D). On the left we show an aggregate having attachment strength $K_a = 1$ dyne/mm, and on the right we show an aggregate having $K_a = 25$ dyne/mm. In these animations, we have ingnored the cells that are in the wetting layer, in line with our analysis presented in Fig. 2C.

#### Movie S5. Comparison of a fluid-like and solid-like migrating aggregate

On the left we show an animation of a fluid-like migrating aggregate corresponding to Fig. 6B. On the right, we show an example of a solid-like migrating aggregate having an average duration of attachment time $T_a = 1$s, whereas the aggregate on the left has $T_a = 10$s. All other parameters are identical for the two simulations.

#### Movie S6. Splitting of a migrating cell aggregate

Here we show the splitting of a migrating cell aggregate, corresponding to Fig. 6C.
